# Novel nuclear role of HDAC6 in prognosis and therapeutic target for colorectal cancer

**DOI:** 10.1101/2020.11.02.356121

**Authors:** Daniel J. García-Domínguez, Lourdes Hontecillas-Prieto, Maciej Kaliszczak, Miaomiao He, Miguel Angel Burguillos, Rajaa Bekay, Vahitha B. Abdul-Salam, Combiz Khozoie, Khalid Shah, Kevin O’Neill, Enrique de Álava, Andrew Silver, Nelofer Syed, Eric O. Aboagye, Nabil Hajji

## Abstract

Histone deacetylase 6 (HDAC6) inhibition is a potential treatment of a wide range of cancer types *via* the acetylation of diverse proteins in the cytoplasm. However, the regulation of histone acetylation and the maintenance of higher-order chromatin structure remains unidentified. Here, we investigated the effect of selective inhibition of HDAC6 by histone acetylation, chromatin relaxation assays, co-immunoprecipitation, acetylome peptide array and *in vivo* RNA microarray. Our data shows that nuclear HDAC6 physically interacts with the Histone 4 lysine 12 residue, and that HDAC6 inhibition increases acetylation specifically at this residue in several cancer types. Inhibition induces major chromatin structure modulation, but has no equivalent effect on knockout HDAC6^-/-^ MEF cells. We identified several novel HDAC6-deacetylated substrates and high expression of HDAC6 in colorectal cancer (CRC) tissue association with reduced levels of H4K12ac and independent of the key CRC driver mutations, but positively associated with *EGFR* expression. Furthermore, *in vivo* HDAC6 inhibition induces significant tumor regression in a CRC xenograft mice model with significant changes in the expression of functional nuclear genes. We also demonstrated that a DNA damaging agent in combination with selective HDAC6 inhibition is effective and acts synergistically, inducing chromatin relaxation and increased cell death in CRC cells. CRC tissues (Normal versus tumor; n=58 matched pairs) together with TCGA data analysis of 467 CRC patients showed that high HDAC6 expression is associated with metastasis, overall and disease-free survival, and is an independent risk factor of CRC stage progression. Our findings designate a new role for nuclear HDAC6 both in cancer prognosis and as a new therapeutic target for CRC and other types of cancer.

**Highlight:** Histone deacetylases 6 activity; Chromatin relaxation; Histone modifications; Gene array; DOX: doxorubicin; OXA: oxaliplatin; 5-FU: fluorouracil; Ac: acetylation; MNase: Micrococal nuclease.

## Introduction

Histone deacetylase 6 (HDAC6) is a member of the class IIb HDAC family and has four function domains: two deacetylase domains, nuclear export sequence (NES), the Glu-containing tetrapeptide (SE14) and the ubiquitin-binding zinc finger domain (ZnF-UBP) (1). HDAC6 is known mainly for deacetylating non-histone substrates. HDAC6 shuttles between the cytoplasm and the nucleus, but predominates in the cytoplasm due to the SE14 motifs, which contribute a strong anchorage of HDAC6 in the cytoplasm (1,2). The major cytoplasmic substrates are VCP/p97, α-tubulin and heat-shock protein 90 (3). HDAC6 activates VCP/p97 ATPase activity to dissociate a heat-shock protein 90/heat-shock factor (Hsp90/HSF1) complex (3). However, no histone lysine residues are known to be deacetylated by HDAC6. HDAC6 inhibition has proved promising as a treatment in several diseases including cancer. Further investigation is now required to shed light on the role of HDAC6 in the cell nucleus. Nuclear HDAC6 has the potential to deacetylate nuclear proteins, such as HDAC11, sumoylated p300, and several transcriptional corepressors including ETO2 (4), and the ligand-dependent nuclear receptor corepressor (LCoR), which is partially nuclear.

HDAC6 is a potential cancer drug target because of its contribution to metastasis via upregulation of cell motility (5). The overexpression of HDAC6 and its correlation with larger tumor size in colorectal cancer (CRC) suggests a key role in CRC progression and drug responsiveness (6). Accordingly, selective HDAC6 inhibition reduces CRC tumor size and inhibits the growth of several colon cancer cell lines (HCT-116, HT29, Caco-2) (6–10). HDAC6 overexpression has also been reported in primary acute myeloid leukemia blasts, in malignant melanoma and human pancreatic cancer tissues (1). Furthermore, the high expression of HDAC6 in estrogen receptor-positive breast cancer increases cell motility by enhancing microtubule activity. Conversely, HDAC6 inhibition decreases microtubule trafficking activity (11). For instance, trafficking of the epidermal growth factor receptor (EGFR) is reduced as it relies on microtubule tracks found to be affected by the loss of HDAC6. Furthermore, EGFR trafficking, found to be modulated by increased α-tubulin acetylation, is mediated by HDAC6 inactivation (12).

The inhibition of HDAC6 has the potential to exert minimal side effects and effectively augment the activity of current anti-tumor drugs (13). Importantly, HDAC6 knockout mouse embryonic fibroblast cells (MEFs) show resistance to transformation and are less prone to cancer (14) whilst selective HDAC6 inhibition is not associated with severe toxicity. For instance, HDAC6 knockout in mice does not lead to embryonic lethality whilst HDAC6-deficient mice are viable and show hyperacetylated tubulin in most tissues (13–20). Nevertheless, HDAC6 inhibitors like Tubacin and Tubastatin A can only be used as experimental tools due to their disadvantageous pharmacokinetic (PK) profiles which have prevented further pre-clinical and clinical development (16, 21). Recently, we developed a novel small molecule hydroxamate (C1A) that preferentially inhibits HDAC6 activity (22). Unlike other HDAC6 inhibitors, C1A has favorable PK *in vivo* and systemic administration of the drug inhibits the growth of colon tumors *in vivo* by up to 78% (22).

To date, the mechanism of selective inhibition of HDAC6 leading to reduced cancer growth has been ascribed mainly to the increased acetylation of cytoplasmic proteins as well as α-tubulin acetylation. Several selective HDAC6 inhibitors have been synthetized, although only Rocilinostat (ACY1215) is used clinically. Its analogue ACY-241 is now available as an oral drug, which is already in Phase 1b clinical development, to be used alone or combined with pomalidomide and dexamethasone in multiple myeloma (NCT02400242)/(NCT02635061). HDAC inhibitors as monotherapy have shown limited success in treating solid tumors (23). However, the combination of HDAC6 with conventional cancer therapies has demonstrated promising anticancer effects in both preclinical and clinical studies (24).

In this present study, we demonstrate that selective inhibition of HDAC6 regulates the acetylation levels of both cytoplasmic and nuclear substrates. Importantly, we show that HDAC6 inhibition increases histone acetylation at H4 lysine 12 (H4K12) as a highly sensitive residue to this inhibition in several cancer types and subsequently induces chromatin relaxation. We also demonstrate that DNA damaging agents in combination with selective HDAC6 inhibition are effective in inducing cancer cell death. We validated the effect of HDAC6 inhibition *in vivo* and verified the level of HDAC6 and its substrate H4K12 in paired patients’ CRC samples (matched normal versus tumor). Furthermore, we validated the importance of high HDAC6 expression in 58 paired primary-metastasis samples and found expression to be associated with CRC metastasis. Importantly, HDAC6 expression was identified as an independent risk factor of CRC progression and associated with a significant reduction in disease-free survival. These findings reveal nuclear HDAC6 role as promising marker for cancer prognosis and as a novel therapeutic target for CRC and other types of cancers.

## Results

### Selective HDAC6 inhibition induces specific histone 4 Lys12 acetylation and chromatin relaxation

The acetylation state of lysine residues regulates both the local chromatin dynamics as well as higher-order chromatin structure. HDAC6 activity has been found to deacetylate several non-histone proteins; however, the inhibition of HDAC6 is not known to regulate the steady-state level of histone acetylation. To explore the effect of selective HDAC6 inhibition on the level of histone acetylation, we predicted the possible interaction with histone tail using the Pathway Commons (http://www.pathwaycommons.org), a collection of publicly available pathways that include biochemical reactions and physical interactions involving proteins and DNA (Figure1, A). The predicted interaction shows mainly the binding to histone 4 family member and, as for nonhistone proteins, HDAC6 was predicted to bind to several EGFR signaling proteins. Furthermore, we predicted the possible interaction of HDAC6 with histone tail residues using All RNA-seq and CHIP-Seq Signature Search Space (ARCHS) that provides access to gene counts from HiSeq 2000, HiSeq 2500 and NextSeq 500 platforms for human and mouse experiments from gene expression omnibus (GEO) and the sequence read archive (SRA) (https://amp.pharm.mssm.edu/archs4/gene/HDAC6) (25). The lysine 12 acetylation in histone 4 tail was the first to be predicted as a substrate for HDAC6 deacetylation activity and the third position among a total gene list with Z-score of 5.03. Overall, this means that H4K12ac is highly predicted by biological processes (GO) to interact with HDAC6 (Figure1, A; Table1). Therefore, we first validated this interaction by investigating the localization of HDAC6 in the nucleus and then subsequently HDAC6’s possible regulation of H4K12 acetylation level. For that, we used the poorly differentiated CRC cell line HCT-116, as we and others have demonstrated that poorly differentiated CRC cells present a low level of H4K12 acetylation compared to highly differentiated CRC cells (26, 27). HCT-116 CRC cells were exposed to 5μM of C1A for 3 hours as a relevant time point that implicates histone acetylation changes in early gene expression regulation (28-30). Both cytoplasmic and nuclear fractions subjected to immunoblotting clearly showed presence of HDAC6 (Figure1, B). Similar results were obtained with BML-281(21) another selective HDAC6 inhibitor (S1, A). No changes in the level of HDAC6 expression were induced by both inhibitors at the concentration of 5μM (Figure1, B and S1, A). Next, we investigated the effect of selective HDAC6 inhibition on H4K12ac and H3K9ac and H4K16ac levels (ranked 3, 21 and 91, respectively; Figure1, A Table1). Using acid histone extraction and western blot analysis, we found that C1A broadly regulates histone acetylation on H4K12 and H3K9 at 3 hours. However, H4K12ac residue showed over sensitivity to HDAC6 inhibition (Figure1, C). H4K12 acetylation was increased in a dose dependent manner starting at the lowest concentration 0.1μM of C1A. In contrast, H3K9 acetylation started to increase at 10 times higher C1A concentration (1μM). C1A treatment did not affect the level of H4K16 acetylation (Figure1, C). In this study, we have shown that the global histone3 and 4 acetylation was not affected by the effect of C1A at 3h exposure. In our previous study, the global histone acetylation even at 24h exposure to C1A 10μM was not affected compared to SAHA (a pan-HDACi) that showed a clear increase in the overall histone acetylation (22).

In order to validate the effect of HDAC6 inhibition, the changes to H4K12ac levels were examined in different cancer types by exposing MCF7 (differentiated breast cancer) (31), U87 (poorly differentiated glioblastoma) (32), and RH30 (poorly differentiated rhabdomyosarcoma) (33) cancer cell lines to increasing concentrations of C1A (0.1, 1, 5 and 10μM). The results showed increased H4K12ac in a dose dependent manner in all the cancer cell lines used (Figure1, D). Furthermore, BML-281 was applied to validate the effect of C1A on the H4K12 acetylation. Both C1A and BML-281 induced similar dose-dependent increases in H4K12ac on the HCT-116 cells (S1, B). The effect of HDAC6 inhibition on the level of H4K12ac regulation was extended by examination in the three cell lines used above and 4 additional cancer cell lines HCT-116 (poorly differentiated CRC), MDA-MB-231 (poorly differentiated breast cancer) (34, 35), CADO-ES (differentiated Ewing sarcoma) (36), and PC3 (differentiated prostate cancer cell) (37) treated with increasing concentrations of BML-281. Similar results as C1A were obtained with BML-281 in all cancer cell lines used (S1, C).

**Figure 1:**
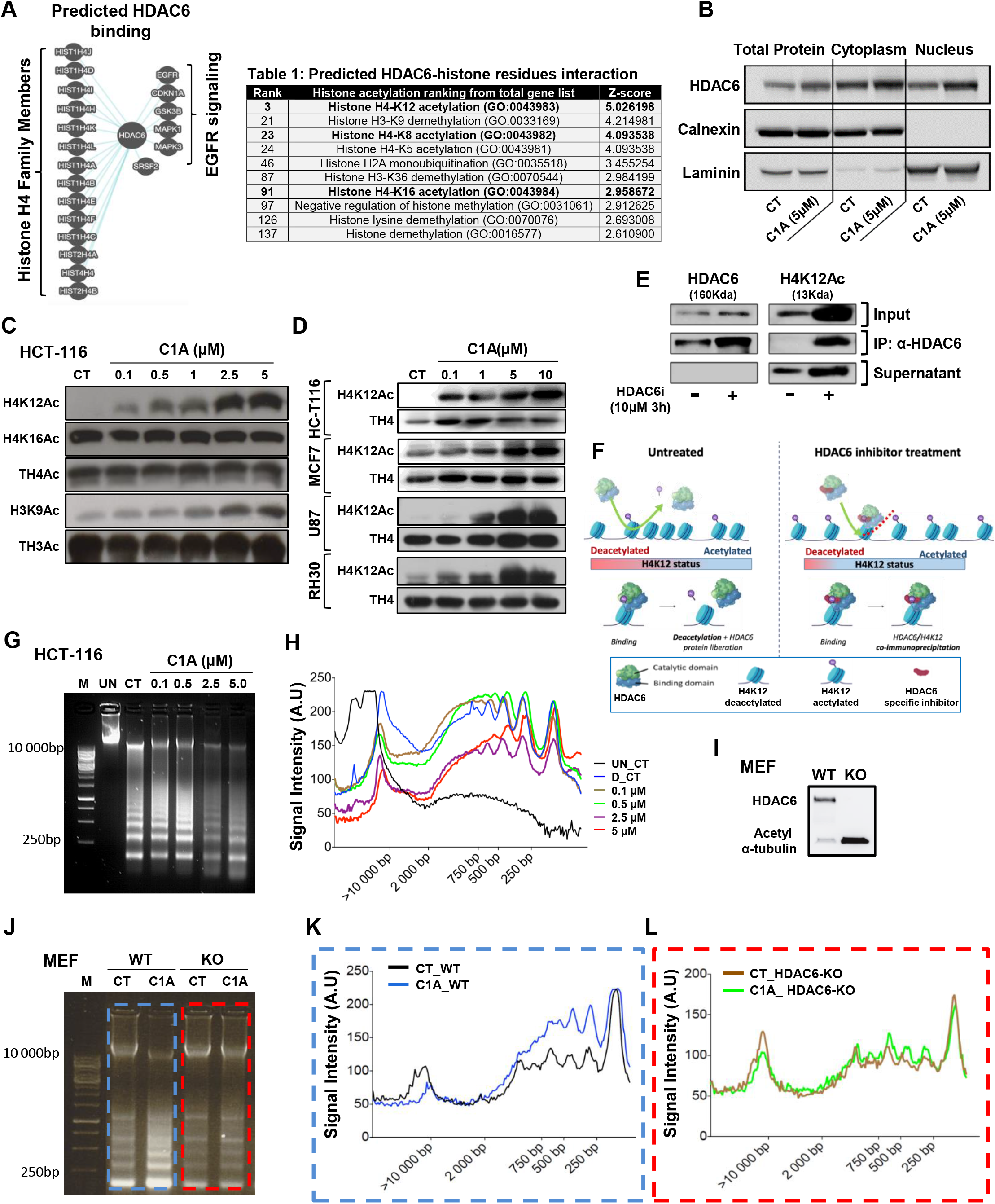
The impact of HDAC6 inhibition on histone acetylation and chromatin relaxation. Pathways common to prediction of HDAC6 binding to histone and non-histone proteins (A). ARCHS prediction of HDAC6 binding to histone lysine residues (Table1). Immunoblotting identification of HDAC6 localization in both cytoplasmic and nuclear fractions (B). Dosedependent evaluation of acetylated level on total H4, H4K12, H4K16, total H3 and H3K9 (C). Immunoblotting evaluation of HDAC6 inhibition and effect on the H4K12ac level in MCF7, U87, and RH30 cancer cells treated with increasing concentrations of C1A (0.1, 1, 5 and 10μM) (D). Assessment of HDAC6 binding to H4 lysine12 residue; Co-immunoprecipitation using the immunoblotting and schematic representation of HDC6-H4K12ac interaction (E and F). Assessment of the impact of HDAC6 inhibition on the level of chromatin structure relaxation by MNase accessibility assay (G) and quantification (H). Western blotting assessment of the level of HDAC6 and acetylated α-tubulin in wild type MEF versus knockout HDAC6 MEF cells (I). Chromatin relaxation assessment by MNase accessibility assay (J). Chromatin relaxation quantification using ImageJ software (NIH) (K and L).

HDAC6 protein contains distinct binding and catalytic domains required for substrates binding and deacetylation (38). Therefore, we explored the possible binding of HDAC6 to the H4K12 residue by co-immunoprecipitation (Co-IP). Cells were treated with HDAC6 inhibitor and Co-IP was performed by using the HDAC6 antibody. The Co-IP results provide evidence of both binding and catalytic action of HDAC6 on the H4K12. While, selective HDAC6 inhibition increased the acetylation on lysine12 residue by inhibiting the function of HDAC6 catalytic domain, the HDAC6 binding domain remain attached to histone 4 as shown by Co-IP (Figure1, E and F).

Since the histone acetylation regulates chromatin higher order structure, we examined the effect of HDAC6 inhibition by C1A on chromatin structure relaxation using the micrococcal nuclease enzyme (MNase) accessibility assay. HCT-116 cells were treated with increasing C1A concentrations and the accessibility assay was performed. The results showed dose dependent chromatin relaxation starting with lowest C1A concentrations; the maximum relaxation was reached at 5μM (Figure1, G and H).

Since we demonstrated recently that C1A could phenocopy HDAC6 genetic knockdown (12), we explored the impact of HDAC6 deficiency in chromatin relaxation. Therefore, we used both mouse embryonic fibroblast wild type (HDAC6 WT-MEF) and isogenic HDAC6 knockout cells with a disrupted catalytic deacetylation domain (12) (Figure1, I). Cells were treated with low concentration C1A (0.5μM) for 3 hours and the MNase assay was performed to assess chromatin relaxation. The results for the Chromatin relaxation assessment showed that the selective HDAC6 inhibition had no effect on chromatin relaxation in knockout HDAC6 MEF cells compared to HDAC6 WT-MEF cells (Figure1, J-L). The use of 0.5μM C1A concentration affect exclusively H4K12 rather than other residues. In exploring the effect of chromatin relaxation this confirms that HDAC6 has a predominant role in the regulation of higher chromatin structure. This inhibition could impact the chromatin status albeit in the initiation of chromatin relaxation which may implicate other residues later.

### Inhibitory effect of HDAC6 at early time-point has no effect on cellular stress

HDAC6 overexpression promotes colorectal tumor growth and selective inhibition synergistically induces CRC cell death (39). Recently we described a selective hydroxamate-based small-molecule HDAC6 inhibitor (C1A), with good pharmacokinetics and ability to modulate autophagy *in vitro* and *in vivo* in different cancer subtypes including CRC cells (12). The mechanism advanced for induction of cell death by C1A, was a mixture of apoptosis and autophagy. Previously, we assessed cancer cell death as a residual effect induced by C1A after 24 hours exposure treatment. Since, the specific epigenetic changes shown in our results above occur at early time exposure (3 hours) to HDAC6 inhibition, we explored whether those changes are a consequence of C1A-induced cell stress at this earlier time-point after C1A exposure. Therefore, ROS generation and DNA damage, as possible cell death triggers, were measured at the early time point. Our data showed very limited levels of ROS generation even with the highest concentration of C1A (10μM); this minor increase in ROS was prevented by antioxidant NAC pretreatment (Figure2, A). Interestingly, the comet assay showed no effectual DNA damage induced by C1A compared to the DNA damaging agent Doxorubicin (positive control) (Figure2, B). The non-appearance of ROS generation and DNA damage with short term C1A exposure even at high concentration indicates that HDAC6 inhibition does not involve cellular stress in the regulation of global acetylation and H4K12ac level or chromatin relaxation as shown in our results (Figure1).

**Figure 2:**
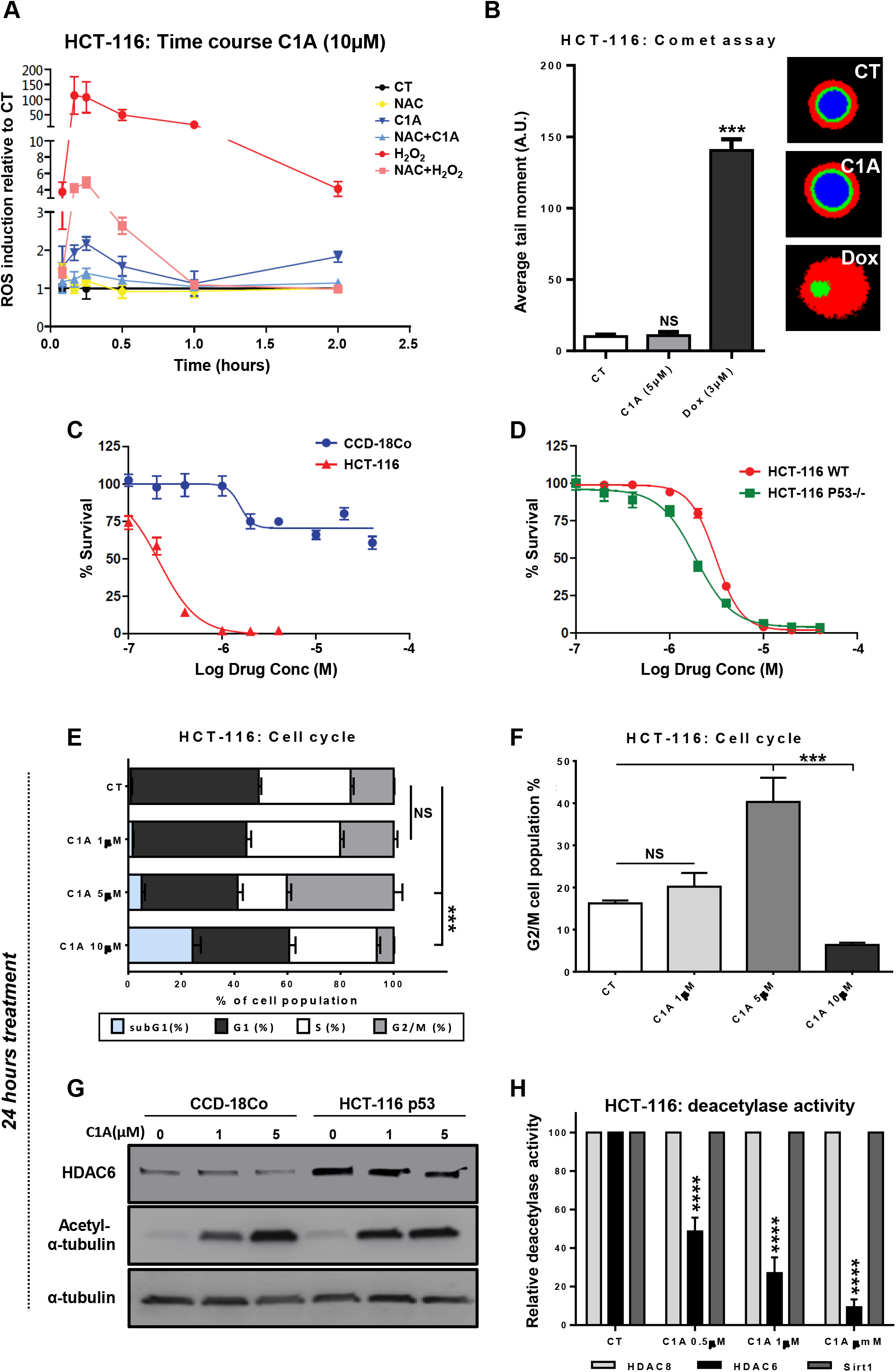
Inhibitory effect of HDAC6 at early time-point has no effect on cancer cellular stress but reduces cellular survival at long-term exposure. Early time course assessment of ROS generation in HCT-116 cells exposed to C1A (10μM) (A). DNA damage evaluation by comet assay in HCT-116 cells treated with 5μM of C1A for 3h (B). Assessment of the effect of varying dose of C1A on cell survival of colonic fibroblasts, CCD-18Co and epithelial cancer cells, HCT 116 p53 and HCT 116 p53^-/-^, using SRB (C and D). HCT-116 cell cycle and cell death analysis by flow cytometry of the effect of varying dose of C1A treatment for 24 h (E-F). Measurement of HDAC6 activity inhibition by acetylated-α-tubulin level (G). Relative deacetylase activity of HDAC6 compared to control HDACs (Sirt1, HDAC8) assessed by immunoprecipitation and *in vitro* HDAC assay (H).

The long-term effect of C1A was explored in dose dependent manner in CRC tumor HCT 116 cells and compared to a different colonic cell type namely fibroblasts from the normal colon CCD-18Co cell line. Our data showed a clear selective sensitivity of epithelial cancer cells compared to fibroblasts cells exposed to C1A for 72 hours. Therefore, C1A significantly reduces HCT-116 cell survival; in contrast, CCD-18Co survived C1A treatment at high concentrations (Figure2, C). In order to evaluate the sensitivity of different CRC cells to C1A, we assessed the importance of p53 status by using wild type and null p53 HCT-116 CRC cell lines. Interestingly, 72 hours exposure to C1A showed dose-dependent results and no statistical differences in percentage surviving cells between wild type and null p53 CRC cell lines (Figure2, D). Furthermore, the CRC cells exposed to different dose-levels of C1A presented significant cell death after 24 hours at 10μM, although cell cycle G2/M arrest was observed only at 5μM of C1A (Figure2, E-F). To confirm HDAC6 activity inhibition both CCD-18Co cells and HCT-116 cancer cells were exposed to 1 and 5μM C1A for 24 hours and the level of acetyl-α-tubulin was measured by immunoblotting. C1A concentrations induced significant increases in the level of acetylated α-tubulin in both cell types (Figure2, G). While, CCD-18Co cells exhibit lower expression of HDAC6 compared to cancer cells, both cell types showed no change in the level of HDAC6 with C1A treatment. Furthermore, the inhibition of HDAC6 activity was measured after 24 hours C1A exposure by immunoprecipitation (HDAC6 enrichment) and *in vitro* HDAC activity assay. Hydroxamic acid derivatives inhibit HDAC6 and HDAC8 with antiproliferative activity in cancer cell lines (40). However, in our previous work we showed that hydroxamate (C1A) preferentially inhibits HDAC6 activity (22). Therefore, we explored the effect of C1A on the HDAC6 and HDAC8 activities. Our results showed that HDAC6 activity was reduced at 0.5, 1, and 5μM, but not the activity of HDAC8 or Sirt1, another well-known to be affected by Hydroxamic acid and used as a control HDACs (Figure2, H).

### Acetylome array identification of novel HDAC6 acetylated markers by MS/MS and UHPLC Q-TOF analysis

The selective inhibition of HDAC6 affects the acetylation level of both α-tubulin in cytoplasm and H4K12 in the nucleus. In order to identify novel acetylated markers targeted by HDAC6 activity, we used the acetylome array in an unbiased analysis to examine possible substrates in both cytoplasm and nucleus fraction. For that, HCT-116 cells were treated with C1A (10μM for 3 hours) and subsequently subjected to detailed characterization by tandem MS/MS and UHPLC/Q-TOF Premier MS analysis.

The HDAC6 substrates were found to be both cytoplasmic (e.g. Tubulin, or heat shock 90kDa protein) and nuclear (e.g. Histone 4, Zing finger protein and histone acetyltransferase Tip 60 protein 3) (Figure3, A and S-Tables1 and 2). All acetylated substrates obtained by acetylome assay were placed into the Global ENRICHNetwork using networkanalyst (networkanalyst.ca) (Figure3, B). As expected, HDAC6 inhibition increases the acetylation of cytoskeleton, microtubules and axon. However, a major effect of HDAC6 inhibition was found to be acetylation of nuclear and cytoplasmic substrates (Figure3, B). Our acetylome data are supported by previous a study where the acetylome was examined using HDAC6 knockout MEF cells. The data showed both nuclear and cytoplasmic lysine deacetylases (KDACs) hyperacetylation (41). Furthermore, the study also explored the overlap between acetylation sites upregulated in cells treated with tubacin (HDAC6 inhibitor) and other HDAC inhibitors. In this previous study with tubacin and our acetylome data with C1A, the use of GO cellular compartment (GOCC) terms for subcellular distribution of proteins with KDACI-upregulated acetylation sites confirm the effect of HDAC6 inhibition of both nuclear and cytoplasmic substrates KDACs.

**Figure 3:**
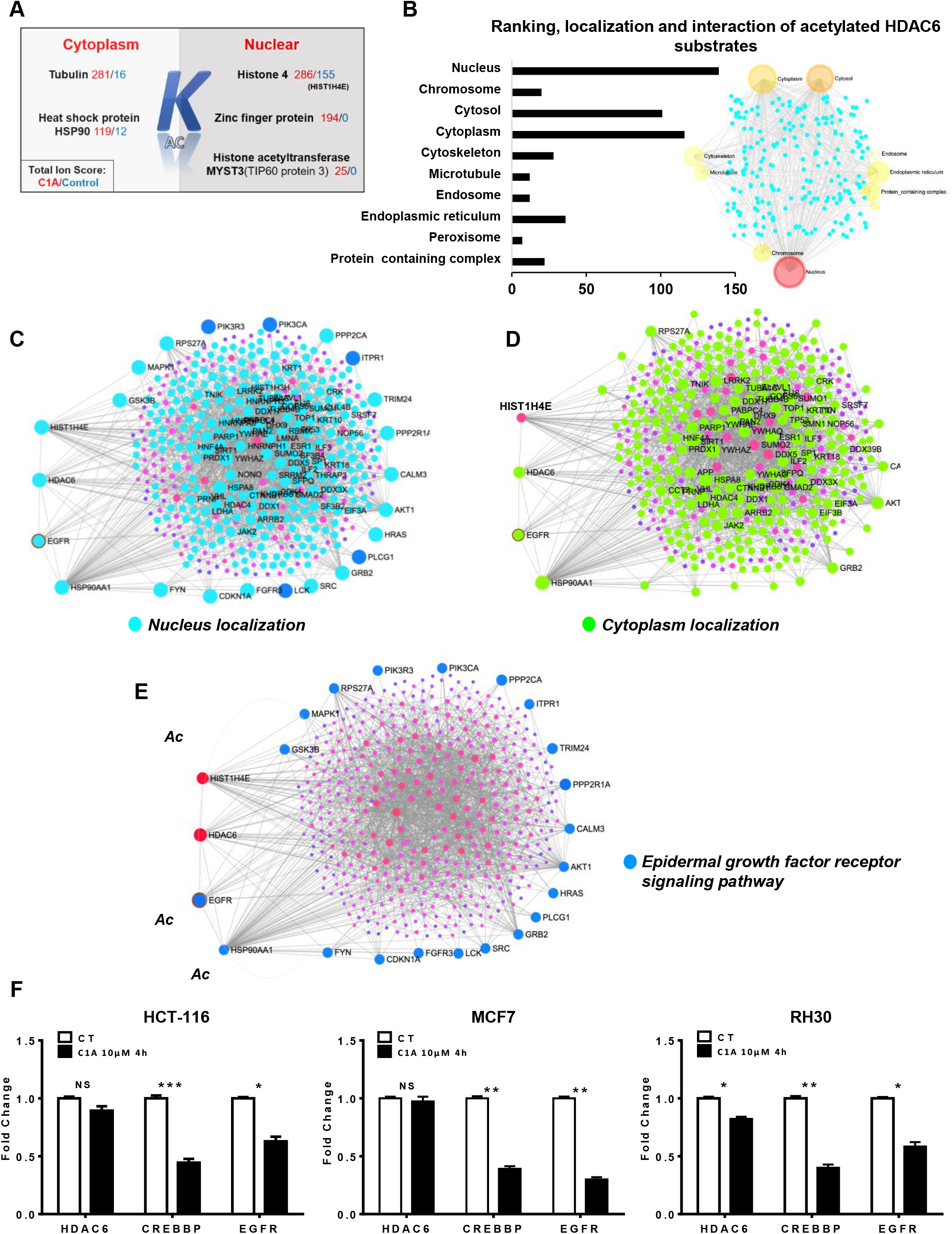
Identification of lysine-acetylated Histone proteins and non-histone proteins in CRC cells. Representative example of total ion score and cellular localization of the peptide after HDAC6 inhibition (A). Assessment of identified protein’s localization obtained by acetylome assay using the Global EnrichNetwork/networkanalyst (networkanalyst.ca) (B). Identification of HDAC6–HIST1H4E interaction networks using the HDAC6 substrates list and uploaded into a visual network-based platform networkanalyst (C-D). Assessment of signaling pathway of the top 20 substrates that interact with HDAC6–HIST1H4E extracted by the ntAct database system (https://www.ebi.ac.uk/intact) and presented by networkanalyst (E). QPCR measurement of *EGFR, CREBBP and* HDAC6 expression level *in* HCT-116, treated with C1A for 3 hours (F). For all the analyses, * p<0.05; ** p<0.01; *** p<0.001 and not significant (ns).

The HDAC6 substrates list (S-Table1, significant hits) was uploaded into a visual networkbased platform networkanalyst to identify HDAC6–HIST1H4E interaction networks. Using the cellular component localization, we identified that the majority of HDAC6 substrates have nuclear and cytoplasmic localization (Figure3, C and D), all of which confirm the subcellular localization of HDAC6 substrates in both cytoplasm and nucleus. Furthermore, we uploaded the top 19 substrates that interact with HDAC6–HIST1H4E obtained by the ntAct database system (https://www.ebi.ac.uk/intact) and presented by networkanalyst (S-Table2). Some interactions were found to be involved in the EGFR signaling pathway (Figure3, E) validating the HDAC6-EGFR interaction as predicted in Figure1A. EGFR acetylation contributes to cancer cell resistance to histone deacetylase inhibitors (HDACis) (42). Moreover, EGFR can be acetylated by CREB binding protein (CBP) acetyltransferase (CREBBP) and the reduction of expression of CREBBP sensitizes cancer cells to HDAC inhibitors (42). In the acetylome analysis, HDAC6 activity was found to deacetylase EGFR and many other substrates in the EGFR signaling pathway. Interestingly, we also predicted that CREBBP is the only protein that connects the HDAC6–HIST1H4E to the EGFR signaling pathway as analyzed by networkanalyst using reactome data base (S2, A).

As global acetylation can regulate gene expression, the effects of HDAC6 inhibition on the expression levels of both *EGFR* and *CREBBP* together with the HDAC6 expression level at an early time point was explored. For that, different cancer types (HCT-116, MCF7 and RH30) were treated with C1A or BML-281 for 3 hours exposure and the mRNA expression level of *HDAC6, CREBBP* and *EGFR* was evaluated by QPCR. Both selective HDAC6 inhibitors (C1A and BML-281) induced significant repression of *EGFR* and *CREBBP* genes in all cancer cell types (Figure3, F and S2, B).

EGFR expression is known to increase proto-oncogene expression levels through caveolin-1 (*CAV1*) activation and cause subsequent cancer cell proliferation. In addition, because pan-HDACis such as TSA and CADC-101 suppress the transcription of EGFR (43–45), we explored the effect of dose dependent selective inhibition of HDAC6 on the expression of HDAC6, *CAV1* and *EGFR* in HCT-116 cells. Both *EGFR* and *CAV1* deceased with early exposure to increasing HDAC6 inhibition, (S2, C). With respect to expression regulation, both EGFR and CAV1 were found to be acetylated by the inhibition of HDAC6 in the acetylome array data (S-Table1, significant hits).

### HDAC6 expression and H4K12 acetylation levels are altered jointly in colorectal cancer patients

To address whether HDAC6 expression is relevant to CRC, we assessed its expression level in two different patient samples cohorts. First, we explored the level of HDAC6 as well as acetylated H4K12 by immunoblotting in an initial series of seven normal-tumor paired CRC patient tissues (Series 1: S-Table3). Our data showed significant overexpression of HDAC6 in CRC patient’s tumor tissues compared to normal tissues. This overexpression mirrored the downregulation of acetylated H4K12 in tumor tissues (Figure4, A-C). Next, we expanded the number of patient’s tissue samples by using a second, independent series for analysis (n=51 normal-tumor paired tissues) (S-Table4). We confirmed by IHC that the positive staining of HDAC6 in colorectal tissues was significantly higher than adjacent normal tissues (Figure4, D and E) in accord with findings for Series 1. The same result was also observed both in glioblastoma and rhabdomyosarcoma (S3, A and B). To investigate the association between HDAC6 expression and CRC patient prognosis and survival, we first assessed the overall survival (OS) and disease-free survival (DFS) by Kaplan Meier analysis (Log Rank - Mantel-Cox test). Our results showed that the differential expression of HDAC6 in CRC patients has a positive correlation on the OS, although this was not significant (*p*=0.0812) (S3, C); but was significantly associated with DFS (*p*=0.0320) (Figure4, F).

**Figure 4:**
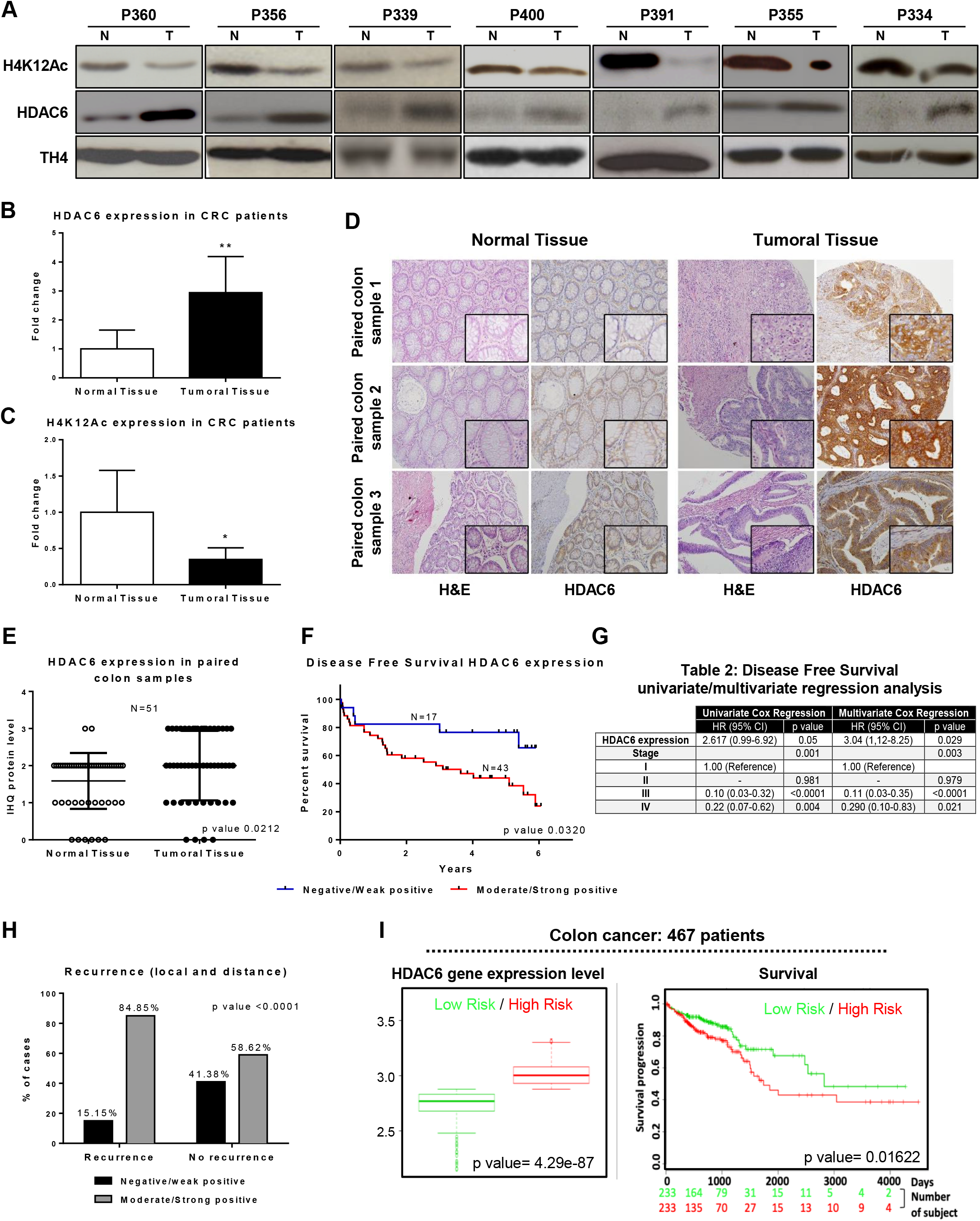
Assessment of HDAC6 expression and H4K12 acetylation levels in colorectal cancer patients. Immunoblotting of samples from colorectal tumor (T) and adjacent normal (N) tissues of 7 paired patients tissue samples were immunoblotted with HDAC6 and H4K12ac antibodies (A). Average quantification of HDAC6 expression and H4K12ac level in the paired CRC versus adjacent normal tissues (B and C). HDAC6 expression and hematoxylin-eosin immunohistochemical staining in representative paired colorectal samples from tissue microarray (20X and 40X magnification) (D). Differential HDAC6 expression between normal and tumoral tissue in 51 paired samples by immunohistochemistry (E). Kaplan-Meier plot of disease-free survival according to the IHC HDAC6 expression in tumoral tissues (F). Table2: Cox-regression analysis of HDAC6 expression and stage in colorectal tumor patients (G). Evaluation of HDAC6 expression levels in local versus distant of tumor recurrence (H). TCGA data analysis of risk factor correlation with HDAC6 expression in colorectal cancer using survExpress website “http://bioinformatica.mty.itesm.mx/SurvExpress” (I, left). Colorectal patient’s survival curves, high risk in red indicates reduced survival; the lower risk group survival is denoted by green curve (I, right). P values calculated using log rank indicate significance at the 95 % confidence level (p < 0.01). Survival analysis was censored by survival days. For all the analyses, * p<0.05; ** p<0.01; *** p<0.001; ****p<0.0001. NS, not significant.

We also examined the prognostic role of HDAC6 expression by Cox-regression analysis of DFS. Here, the clinical stage at diagnosis (a critical prognosis factor in CRC patients) and the expression of HDAC6 was explored in DFS. Univariate and multivariate analysis showed that HDAC6 expression was strongly associated with DFS in a cancer stage-independent manner (*p*=0.05 and *p*=0.029, respectively (Figure4, G; Table2). Additionally, CRC clinical stages correlating with DFS (Table2) were in agreement with the Kaplan–Meier analysis (S3, D). These results showed that HDAC6 expression and the clinical stage at diagnosis were both independent prognostic indicators of DFS. Since HDAC6 expression correlated with DFS, we also evaluated whether tumor HDAC6 levels were associated with tumor recurrence (local and distant). Interestingly, analysis of Series 2 revealed that the percentage of recurrence cases were significantly higher in tumors with high HDAC6 expression (85%) than with low HDAC6 expression (15%) (Figure4, H), which is in line with our results. As a further validation, TCGA data analysis of 467 CRC patients revealed that HDAC6 overexpression correlated with high risk of death and significant reduction in survival (Figure4, I). Moreover, results comparable to those obtained in CRC were found in breast cancer, ovarian and sarcomas in which the positive HDAC6 transcript expression achieved a significant high risk in patients with high HDAC6 expression (S3, E).

It has been suggested that the metastatic CRC resistance mechanisms are not linked with the occurrence of *K-Ras* and *BRAF* mutations or quantitative change in the *EGFR* expression (46–48). Based on the acetylome and RT-qPCR analyses, we propose that *HDAC6* expression is connected to *EGFR* through the regulation of both acetylation and expression. In order to explore the relationship between *HDAC6* and *EGFR* in wider numbers of cancers, we analyzed TCGA data base for both the correlation between the expression of *EGFR* and *HDAC6*, as well as the association of mutated *EGFR* and *HDAC6* expression in 40 types of cancers (S-Table5, S-Table6, and S4, A and B). The results demonstrate a clear absence of association between mutated EGFR and HDAC6 expression, with only 2.5% of 40 cancer types (i.e. SKCM-primary) showing a positive relation and 5% a negative relationship (i.e. HPV-Positive and negative HNSCC) (S-Table6, and S4, B). While 92% of cancer showed no association of *HDAC6* expression level with mutated *EGFR* including CRC, 67.5% of cancers showed a positive association of EGFR and *HDAC6* expressions including in CRC (S-Table6 and S4, A and B). Interestingly, *KRAS, HRAS, ARAF* and *BRAF* mutations showed no association with *HDAC6* expression in all 40 types of cancers including CRC with exception of SKCM-primary cancer that shows only association of *HRAS* mutation with HDAC6 expression (S-Table7, S4 C-F). This suggested that *EGFR* and *HDAC6* expression are connected positively, but are independent of these mutations in tumors.

### C1A-treated xenografts in a mice model induces tumor regression and significant reduction of HDAC6 at mRNA and protein levels

In order to substantiate the effect of C1A on tumor growth, female Balb-c nude/nude mice were injected subcutaneously with HCT-116 cells. A significant reduction in tumor growth was detected from day 7 (Figure5, A). The *in vivo* experiments were terminated on day 14, where significant reduction in the tumors size was observed in C1A treated animals (Figure5, B). Then, tumors were excised and total protein and mRNA extracted and analyzed. Significant decrease of HDAC6 was found at both protein and mRNA levels of pooled samples as shown by immunoblotting and gene microarray (Figure5, C and D). Furthermore, *in vivo* gene array analysis of isolated tumor from HCT-116 xenograft mice showed significant regulation of the transcription of several genes (S-Table8). The affected genes have regulative role essentially in the nucleus, cytoplasm and cytoskeleton as represented Global EnrichNetwork/networkanalyst (Figure5, E and S-Table8).

**Figure 5:**
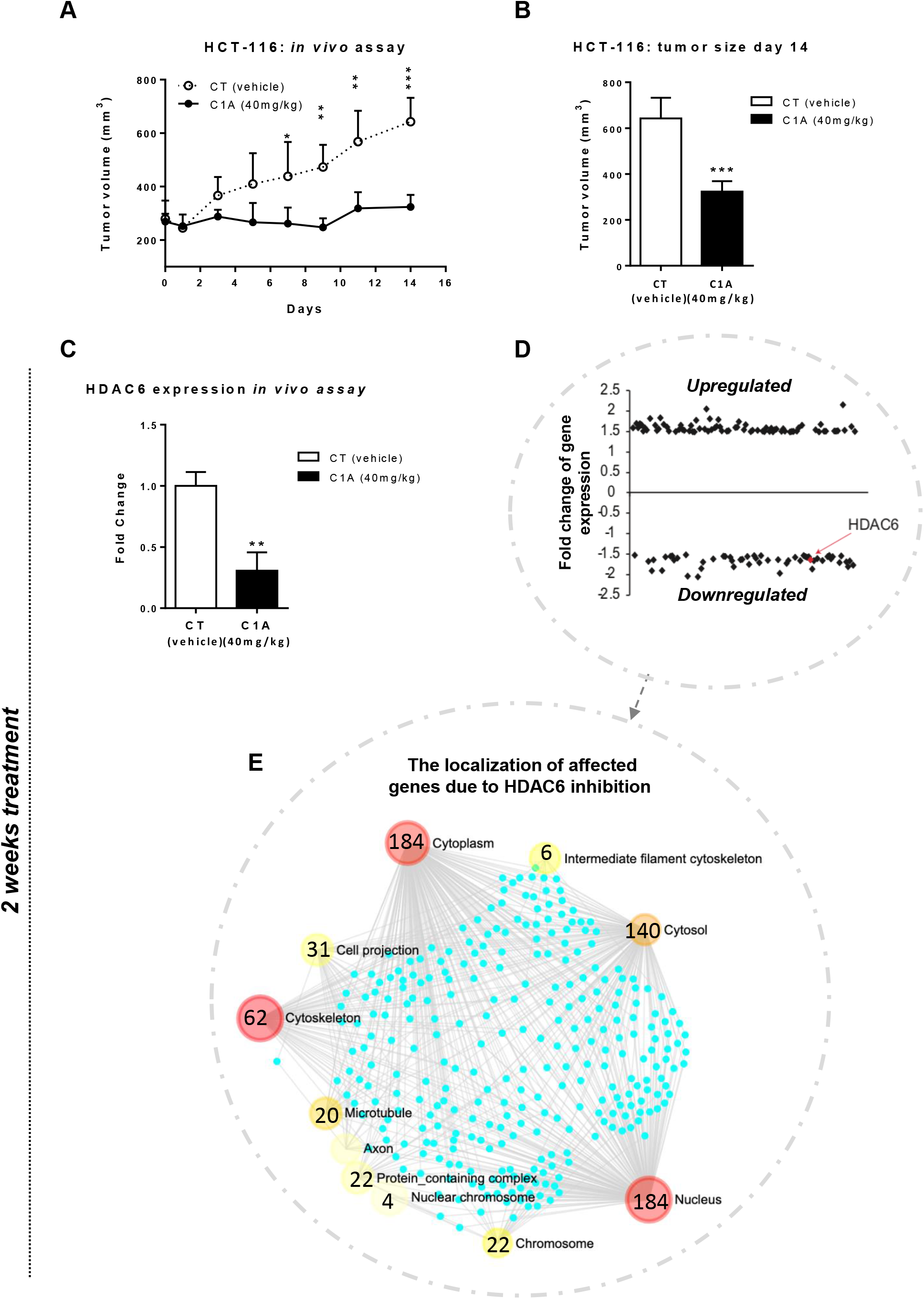
*In vivo* analysis of the effect of selective HDAC6 inhibition. Xenograft HCT-116 mice model treated with C1A for 14 days. Average tumor volume assessment was made before treatment (day 0) and every two days during the study using calipers (A). Histogram representing the average of tumor volume on day 14 (B). Tumors were excised on day 14 and analyzed for HDAC6 protein level by immunoblotting for groups of 5 animals (C). Representation of regulated genes after 2 weeks in mice treated with C1A (D). Network analysis of *in vivo* C1A affected genes interaction and cellular localization from C1A treatment tumors (E). For all the analyses, * p<0.05; ** p<0.01; *** p<0.001. NS, not significant.

### Combined treatment with C1A and DNA damaging agents induces a significant increase of H4K12ac, acetylated a-tubulin, chromatin relaxation and subsequently enhances significant CRC cell death

Next, we investigated why the low levels C1A do not induce a direct cell kill, but modifies H4K12ac and then later enhances cancer cells sensitivity to conventional treatment. First, we explored the effect of individual potencies of DNA damaging agents by treating HCT-116 cells with selected concentrations (low, medium and high) of doxorubicin (Dox); oxaliplatin (Oxa) and fluorouracil (5-FU) for 24 hours. After treatment, the cells were stained with DiOC6(3) and propidium iodide (Pi) for measurement of mitochondrial membrane potentials (Δψm) and cell death, respectively. Mitochondrial membrane permeabilization (MMP)-inducing agents lead to mitochondrial transmembrane potential (Δψm) loss and subsequent release of soluble intermembrane proteins, which are the signals of apoptosis. Our data show that the loss of Δψm and cell death are increased in a dose-dependent manner (Figure6, A and B). To determine apoptotic cell death pathway induced by DNA damaging agents, we next investigated the level of PARPc as an apoptotic hallmark, P53 stabilization as a pro-apoptotic protein and survivin expression as an apoptosis inhibitor. Our data shows clear dose-dependent modulation of PARPc increase, P53 stabilization and downregulation of survivin (Figure6, C). Recently, we reported that SAHA (vorinostat) as a pan-HDAC inhibitor known to inhibit HDAC6 activity and also other HDACs (49), augments the effect of DNA damaging agents both *in vitro* and *in vivo*. Importantly, we found previously that SAHA not VPA (classI and IIa inhibitor) as monotherapy or combined with DNA damaging agents induces significant increases in H4K12 acetylation in several CRC cell lines (i.e. HCT116 p53+/+, HCT116 p53-/-, SW480, HT-29) and subsequently induces significant chromatin relaxation (27).

**Figure 6:**
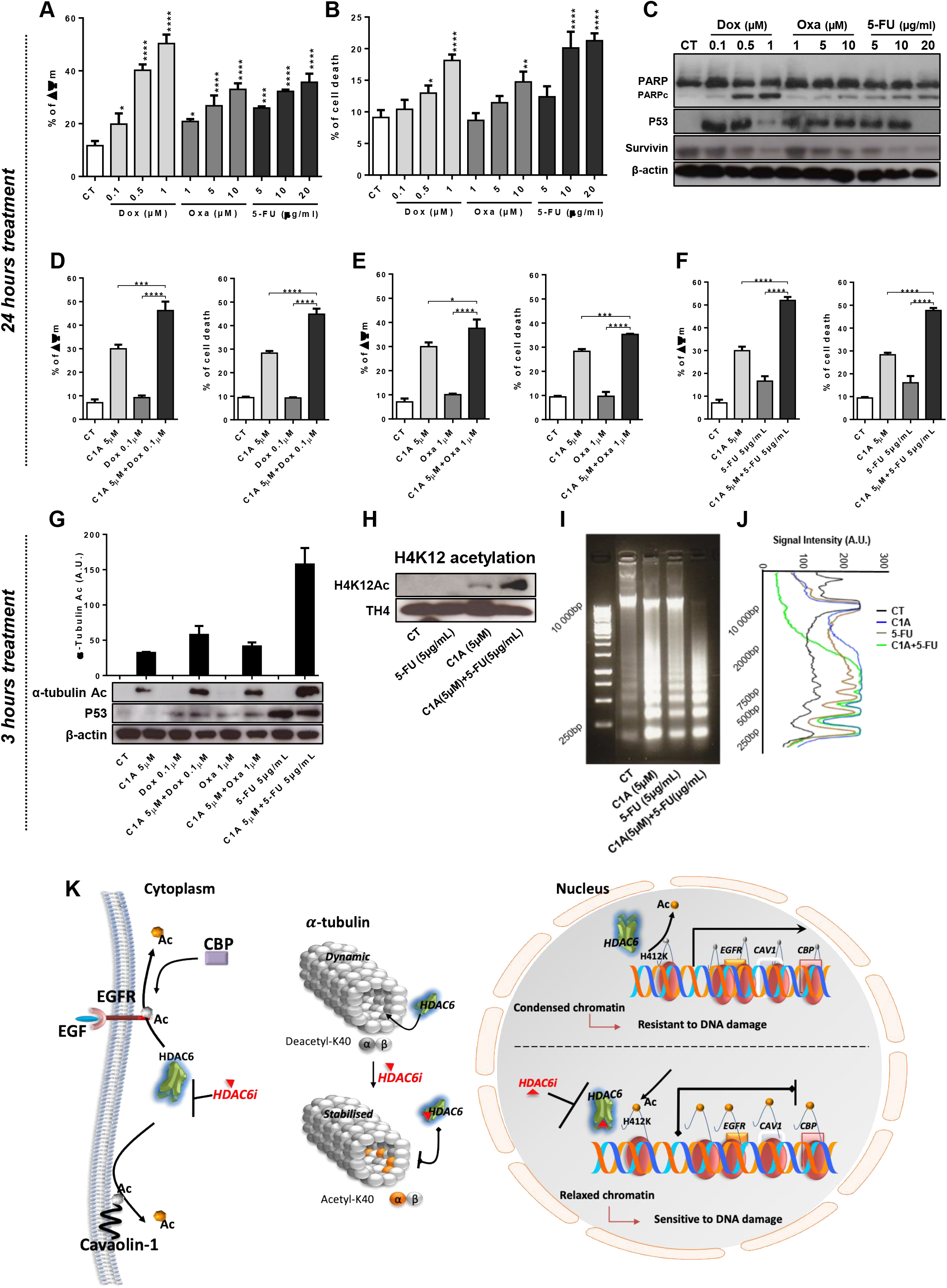
Assessment of the combined treatment C1A and DNA damaging agents. Mitochondrial potential loss (Δψm) assessment in HCT-116 cells exposed to varying drug concentrations for 24 hours stained with DiOC6(3) and analyzed by flow cytometry (A). Cell death (Sub-G1) assessment of the combined treatment by flow cytometry (B). Immunoblotting for evaluation of PARPc (apoptotic hallmark), P53 protein and survivin levels (C). Assessment of the combined treatment C1A and DNA damaging agents on the levels mitochondrial potential loss (Δψm) and cell death (Sub-G1) (D-F). Immunoblotting evaluation of 3 hours effect of combined treatments on the level of acetylated α-tubulin (G). Assessment of 3 hours combined treatment C1A/5-FU on the levels of H4K12ac, as well as chromatin structure relaxation (I-J). Chromatin structure relaxation quantification using ImageJ software (NIH) (J). Schematic representation of HDAC6 deacetylation activity by C1A in cytoplasm and nucleus (K). For all the analyses, * p<0.05; ** p<0.01; *** p<0.001; **** p<0.0001. NS, not significant.

Accordingly, the combined effect of C1A and DNA damaging agents was investigated by using the lowest drug concentrations with 5μM C1A for 24 hours. Our results demonstrated that the combined treatment is greater than the individual drugs potencies (Figure6, D-F). Synergistic effects were further identified in the combinations Dox+C1A and 5-FU+C1A, while additive effects were seen in the OXA+C1A combination.

Assessment of the combined treatment effect on the level of HDAC6 activity in deacetylating α-tubulin was assessed after cells were exposed to C1A for a short period (3 hours). While DNA damaging agents as monotherapy have no effect on the level of acetylated α-tubulin, the combination with C1A induces synergistic acetylated α-tubulin with respect to C1A alone (Figure6, G).

The most effective combination was found to be the first-choice chemotherapy drug for CRC, 5-FU, together with C1A. We examined this combination further at the early time point (3 hours) by measuring the level of H4K12 acetylation, as well as chromatin relaxation. Our data show that this combined treatment induces a major increase in both H4K12ac and chromatin relaxation levels (Figure6, H-J).

Our data conclusively identifies a novel role for HDAC6 deacetylating activity: HDAC6 was found to regulate the acetylation level of H4 at lysine12 leading to chromatin relaxation and transcription repression of *EGFR*, *CAV1* and *CREBBP* and to subsequently increase the sensitivity of cancer cells to DNA damaging agents. In addition, to the known classical substrate α-tubulin, HDAC6 was found to deacetylate several cytoplasmic and nuclear substrates including EGFR and CAV1. Both EGFR and CAV1 acetylation and expression levels were shown to be regulated by the selective inhibition of HDAC6. Furthermore, the CERBBP acetyltransferase activity on EGFR acetylation was found to be balanced by HDAC6 activity (Figure6, K).

## Discussion

Histone deacetylases (HDACs) are characterized as nuclear enzymes deacetylating histone tails. However, several non-histone proteins have been identified as substrates of HDACs in both the cytoplasm and nucleus. The subcellular localization of HDACs is considered a key epigenetic factor that regulates the higher order chromatin structure and represses genes expression. HDAC family consists of four classes highly conserved from invertebrates to mammals (50). Class II HDACs are known to shuttle between the nucleus and the cytoplasm with the exception of HDAC6, which is localized mainly in the cytoplasm to deacetylate α-tubulin, Hsp90, and cortactin (51). HDAC6 is a unique class IIb HDAC which contains two deacetylase domains responsible of deacetylation of a number of substrates involved in tumorigenesis, cell survival, cell motility, and transcriptional or translational response (52). Compelling evidence has accumulated to support the hypothesis that HDAC6 deacetylates nuclear substrates and is localized partially in the nucleus to interact with several non-histone nuclear proteins, including HDAC11, the transcriptional corepressor LCoR, and transcription factors such as NF-κB and Runx. Interestingly, HDAC6 was found to be acetylated by p300, and mutations of the lysine(s) responsible of nuclear HDAC6 localization retained HDAC6 in the cytoplasm by blocking the interaction with the nuclear import protein importin-α (53). The above account infers nuclear localization of HDAC6. However, HDAC6 activity is not known to regulate the acetylation level of histone residues.

Here, we first predicted a possible physical interaction of HDAC6 with histone tail residues. Importantly, the bioinformatics prediction showed that the HDAC6 interacts only with histone 4 family members. We further predicted that the lysine 12 residue on histone 4 was a highly sensitive residue that could be affected by the HDAC6 activity. Initially, we validated this prediction by cellular fractionation identifying the presence of HDAC6 in both cytoplasmic and nuclear fraction. Furthermore, the predicted lysine was investigated by histone acid extraction in CRC cells HCT-116 (26).

As predicted, H4K12ac showed high sensitivity to the selective HDAC6 inhibitor C1A at the early time exposure (3 hours) compared to other histone residues. Interestingly, it is known that the poorly differentiated CRC cells showed low levels of H4K12 acetylation compared to the highly differentiated CRC cells (26). The HCT-116 cells used here are poorly differentiated CRC cells that show undetectable levels of acetylation at the H4K12 residue. Importantly, increases of C1A concentrations induces significant gradual increases of acetylated H4K12. We validated this observation on several cancer types exposed to C1A or BML-281 as selective HDAC6 inhibitors. Furthermore, Co-IP results showed that selective inhibition of HDAC6 increases the acetylation on lysine12 residue together with increased HDAC6 binding to H4K12. Interestingly, similar observations were reported previously between the interaction of HDAC6 and microtubules. The selective inhibition of enzymatic activity of HDAC6 by Tubastatin A increases its binding to microtubules. This binding enhanced microtubule dynamic instability in breast cancer cells (54). These findings suggested that HDAC6 might act as a microtubule-associated protein (MAP) regulating microtubule dynamics under certain conditions. Our data could also indicate that HDAC6 binding to H4K12ac may function as chromatin-associated protein controlling its dynamic as a necessary requirement for cancer cell maintenance.

Since chromatin remodeling is tightly linked to the level of histone acetylation, we explored the effect of a gradual increase of H4K12 acetylation induced by C1A dose-dependent on the chromatin relaxation. Our data showed for the first time that selective HDAC6 inhibition induces chromatin relaxation by increasing the level of H4K12ac. These results are in concordance with our previous observation, that Vorinostat or suberoylanilide hydroxamic acid (SAHA), as a pan-HDAC inhibitor and the most widely used inhibitor of class II HDAC activity, induces significant increase in the acetylation in the H4K12ac in CRC cell lines including HCT-116, HCT-116-P53^-/-^, SW480, HT-29 (27). We also found in this study that SAHA induces chromatin relaxation in CRC cells (27) in a similar manner to C1A. Additionally, we showed that the disrupted catalytic deacetylation domain of HDAC6 prevents C1A from inducing of chromatin relaxation. This also confirmed our recent observation that the effect of C1A on CRC cells reproduces a similar signaling pathway mechanism as if induced by genetic knockout of HDAC6 (12).

Critically, the acetylome objectively identified H4 acetylation as a substrate of HDAC6 in C1A treated CRC cells. As expected, several HDAC6 substrates were found to be involved in the regulation of cytoskeleton, microtubules and axon. The acetylome analysis revealed a notable presence of nuclear and cytoplasmic substrates, supporting the hypothesis that there is a nuclear role for HDAC6 localization. The acetylome also validated the bioinformatics prediction of HDAC6 interaction with the EGFR signaling pathway. Moreover, the inhibition of HDAC6 activity reduced both the expression level of *EGFR* acetyltransferase *CREB* binding protein (CBP) and *EGFR* in the different cancer types examined.

EGFR is known to play a crucial role in cancer and normal cell growth, differentiation, and motility (55). EGFR is frequently overexpressed in variety of cancers and it is known that its acetylation contributes to cancer resistance to HDAC inhibitors; the reduction of expression of *CREBBP* has the opposite effect. EGFR expression regulates the increase of proto-oncogenes like *c-fos* expression level through caveolin-1 activation, which then leads to cancer cell proliferation (56). The use of pan-HDACi TSA is known to suppress EGFR transcription and subsequently repress cellular proliferation (57). Similarly, our results showed that the inhibition of HDAC6 using C1A or BML-281 is sufficient to reduce EGFR and *CREBBP*, as well as *CAV1*, expressions. The selective inhibition of HDAC6 emphasized the exclusive role of HDAC6 among other HDACs in regulating EGFR signaling pathway. Furthermore, EGFR status associated with *K-RAS* and *BRAF* mutations, as important markers in CRC, does not provide any relevant information concerning mechanisms of resistance in metastatic CRC and does not guarantee the efficacy of anti-EGFR therapy (58). Our results are in agreement and show that *KRAS, HRAS, ARAF* and *BRAF* mutations have no association with *EGFR* expression in most cancers including CRC. However, *EGFR* and *HDAC6* expression found in our TCGA data analysis were connected positively and are independent of these mutations in tumors.

The overexpression of HDAC6 was found to be associated with larger tumor sizes in CRC indicating its key role in CRC progression (6, 39). Importantly, selective HDAC6 inhibition reduced CRC tumor size and inhibited the growth of several colon cancer cell lines (HCT-116, HT29, Caco-2) (39). The overexpression of HDAC6 was identified in a variety of cancers (1) which suggests a significant role for HDAC6 in cancers including CRC, primary acute myeloid leukemia blasts, malignant melanoma, primary oral squamous cell lines, breast cancer and human pancreatic cancer tissues (1). Furthermore, the high expression of HDAC6 in estrogen receptor-positive breast cancer increases cell motility by enhancing microtubule activity (59).

It has also been shown that the overexpression of HDAC6 and its cytoplasmic deacetylating activity is required for efficient oncogenic cellular transformation (14). Transformed normal human embryonic kidney cells (HEK), and prostate epithelial cells (PrEC) transformed with retrovirus expressing SV40 exhibit high levels of HDAC6 expression driven by oncogenic Ras (1). This indicates further a critical role for HDAC6 in tumorigenesis and supports the hypothesis that the inactivation of HDAC6 in several cancers could be beneficial in reducing tumor growth. Here, we identified a significant overexpression of HDAC6 in CRC tumors accompanied by a significant repression of its substrate H4K12ac compared to normal patients’ tissue. While, the low-level expression of HDAC6 correlated significantly with the improved DFS, the overexpression was also associated significantly with CRC metastasis. The significant correlation between DSF and HDAC6 expression was also independent of other relevant clinical parameters in CRC, such as stage. This positions HDAC6 as a molecular factor with a relevant prognostic value in CRC. Moreover, we showed that the inhibition of HDAC6 activity *in vivo* significantly decreases tumor volume with significant modulation of genes whose expression have critical regulative roles in the nucleus, cytoplasm and cytoskeleton. Long-term *in vivo* C1A treatment was not only beneficial in reducing tumor volume, but also reduced significantly HDAC6 expression at protein and mRNA levels.

C1A was developed based on the structure of SAHA to preserve the hydroxamate part of the molecule that binds to the Zn^2+^ pocket of HDAC. In general, C1A was found to enhance preferential binding to HDAC6 catalytic domain (cd) II (22). C1A induced sustained acetylation of HDAC6 substrates, a-tubulin and HSP90, compared with current clinically FDA approved HDAC pan-inhibitor SAHA. It is known that the inhibition of HDAC6 is not associated with severe toxicity. As with C1A, we showed that SAHA increases significantly the effect of DNA damaging agent DOX. SAHA is widely known to induce very low toxicity (60), although as a pan-HDAC inhibitor it can affect the activity of non-relevant HDACs such as HDAC1 and HDAC2 in cancer. In contrast, C1A is a selective and effective HDAC6 inhibitor. In this previous study, we identified a clear correlation between histone acetylation level and the sensitivity of CRC cell lines to SAHA combined with DNA damaging agents. The levels of histone acetylation in HCT-116 p53+/+, HCT116 p53-/-, SW480, HT-29 were explored and we showed that SAHA as single treatment was much more potent cross all the different CRC cell lines inducing significant increase in the H4K12 acetylation compared to the VPA (classI and IIa HDAC inhibitor) (27). In addition, the combined treatment SAHA+Dox increased significantly the level of H4K12ac compared to the combined VPA+Dox, which showed no significant increase in H4K12ac. This suggest that classI and IIa HDACs are most likely not involved in the regulation of H4K12ac.

The effect of C1A at early time point, as shown by H4K12ac and chromatin relaxation, appears not to involve DNA damage induction as indicated by comet assay and ROS measurement. Nevertheless, HDAC6 inhibition with C1A appeared to selectively eliminate CRC cancer cells, but not normal colon cells. In general, this histone deacetylase inhibitor induces G2/M cell cycle arrest and cancer cell death independently of DNA damage (61, 62). This was found to be in line with long-term CRC exposure to C1A, which also induces cell cycle G2/M arrest followed by cell death. Our results show that HDAC6 inhibition by C1A synergistically augments the efficacy of conventional DNA damaging agent on CRC cells by inducing apoptotic cell death. The possible mechanism identified was associated with a synergistic increase in α-tubulin acetylation, H4K12ac and the level of chromatin relaxation (Figure6, K). We have previously demonstrated that the inhibition of certain HDACs regulates the level of histone acetylation inducing chromatin relaxation and increasing the accessibility of drugs that act on DNA (63). Therefore, C1A, by inhibiting HDAC6, increases the level of H4K12 acetylation leading to chromatin relaxation, which subsequently increases the efficacy of 5-FU to induce cell death probably *via* DNA damage. This mechanism could be beneficial not only for CRC, but for cancers that present low level of H4K12 acetylation as the global hypoacetylation of H4K12 was considered to be informative of CRC progression (26). Here, we determined that HDAC6 regulates this specific residue, which controls the higher order chromatin structure. Altogether, our pre-clinical investigations emphasize the importance of HDAC6 as component in cancer development and the marked antitumor effects of selective inhibition of HDAC6 by selective inhibitors (e.i. C1A and BML281) when used in combination with DNA damaging agents. Further investigations to appropriately combine conventional DNA damaging agents with HDAC6 selective inhibitor should have significant clinical importance and are warranted.

## Supporting information

supplementary data. Dani et al HDAC6

## Acknowledgments

We would like to thank Imperial College department of medicine, Bain division, also to thank the donors and the HUVR-IBiS Biobank (Andalusian Public Health System Biobank and ISCIII-Red de Biobancos PT13/0010/0056) for the human specimens used in this study. EOA would like to thank Cancer Research UK (C2536/A10337 & C2536/A16584), Imperial College Experimental Cancer Medicine’s Centre and Biomedical Research Centre. NH would like also to thank the great support from Brain Tumor Research Campaign (BTRC) and Brain Tumor Research (BTR). DGD and LHP are supported by CIBERONC (CB16/12/00361). LHP funded by the Consejería de Salud, Junta de Andalucía (PI-0013-2018).

## Material and methods

### Cell lines

Colorectal cancer (HCT-116, HCT-116 P53-/-), breast cancer cells (MCF7, MDA-MB-231), glioblastoma (U87), adult sarcoma CADO-ES cells, pediatric sarcoma (rhabdomyosarcoma RH30), prostate cancer (PC3) were used. HCT-116 cells were obtained from the American Type Cell Culture Collection (Rockville, MD, USA) and authenticated by short tandem repeat profiling under contract by DDC Medical (London, UK). HCT-116 cells P53-/- were generously supplied by Prof. Simak Ali (Imperial College London, UK). U87 glioblastoma cells were generously supplied by Prof. Venero Recio (University of Seville, Spain). MCF7 and MDA-MB-231 cells were kindly provided by Dr. Pérez Losada Jesus (Cancer research institute of Salamanca, Spain). PC3 cells were kindly provided by Dr. Sáez (Institute of Biomedicine of Seville, Spain). RH30 and CADO-ES cells were obtained from the EuroBoNet cell line panel. Mouse Embryonic Fibroblasts (MEFS WT mutant and KO in HDAC6) were kindly provided by Prof. Tso-Pang Yao from Duke University School of Medicine (Durham, NC, USA).

All cancer cells were maintained in DMEM medium containing 2mM L-glutamine, except RH30, CADO-ES and PC3 cells were maintained in RPMI medium. Both mediums were supplemented with 1% penicillin/streptomycin and 10% (v/v) fetal calf serum (Sigma-Aldrich, St. Louis, MO, USA) and grown at 37°C, 5% CO2. All cells were free of mycoplasma, as screened with the MycoAlert® Mycoplasma Detection Kit (Lonza).

### HDAC6 inhibitors compounds

C1A (synthesized in-house)(22). BML-281 was purchased from Enzo Life Sciences Inc. (UK). Stock solutions of both compounds were prepared in dimethyl sulfoxide (DMSO) and diluted to final concentration in the culture medium 1:1000 (v/v).

### Subcellular fractionation

Preparation of nuclear and cytosolic cell fractions. Human breast cancer cell lines were harvested at 80% confluence through trypsination. Isolation of nuclei and cytosol was carried out using NE-PER Nuclear and Cytoplasmic Extraction Reagents (Pierce, Bonn, Germany) following the manufacturer’s instructions. Probes were solved in Laemmli sample buffer at a final concentration of 1×10^6^/ml and stored at −20°C before Western Blot electrophoresis. Preparation of nuclear and cytosolic cell fractions. Human breast cancer cell lines were harvested at 80% confluence through trypsination. Isolation of nuclei and cytosol was carried out using NE-PER Nuclear and Cytoplasmic Extraction Reagents (Pierce, Bonn, Germany) following the manufacturer’s instructions. Probes were solved in Laemmli sample buffer at a final concentration of 1×10^6^/ml and stored at −20°C before Western Blot electrophoresis. To isolate sub-cellular fractions, cells were lysated in 800μl of 10mM HEPES pH 7.9 buffer, containing 10mM KCl, 0.1mM EDTA and 0.1mM EGTA on ice. Lysate cells were incubated on ice for 15 minutes, vortex the tubes for 5 seconds and centrifuge the tubes for 2 minutes at 13,000rpm at 4°C. The supernatant (cytoplasmic fraction) was transferred to a clean pre-chilled tube and place this tube on ice until use or storage at −80°C. The nuclear fractions recovered in the pellet were resuspended in 75μl of 20mM HEPES pH 7.9 buffer, containing 0.4M NaCl, 1mM EDTA, 1mM EGTA, and the protease inhibitors cocktail solution. After 15 minutes of incubation on ice and vortex, tubes were centrifuged for 5 minutes at 14,000rpm 4°C. The supernatant (nuclear fraction) was transferred to a clean pre-chilled tube and place this tube on ice until use or storage at −80°C. Reagents: HEPES Applichem (A1069), KCl Applichem (A1362), EDTA Applichem (A3234), EGTA Sigma-Aldrich (E4378), NACL Applichem (A1149).

### Extraction, isolation and purification of histones

Cancer cell lines were harvested at 80% confluence through trypsination. Extraction, isolation and purification of histones were carried out using the protocol described previously(64).

### Co-Immunoprecipitation assay

Whole protein extracts (250μg) in NP40 buffer (150mM NaCl, 20mM Tris pH 8.0, 1mM DTT, 0.5% NP40) were incubated with 15μl of protein A-dynabeads (Invitrogen) coupled to 2μg of HDAC6 Cell Signalling (7558), 3 hours at 4°C. After magnetic immunoprecipitation and washes, immunoprecipitates were resolved in a 10% polyacrilamyde SDS-PAGE gel, transferred and blotted as described above using H4K12ac antibody (ab177793).

### Total protein extract

Cells were cultured for 24h and subsequently treated with different compounds: C1A or BML-281 at the indicated time and concentrations. Protein samples were subsequently prepared by lysing cells in RIPA buffer (Life technologies) supplemented with protease and phosphatase inhibitor cocktails (Sigma) and subject to standard western blot procedures.

### Immunoblotting

Histone, nuclear and cytoplasm protein fractions, or total protein extracts were quantified by Bradford assay. Western blot was performed using standard protocols. Incubation with primary antibodies was done at 4°C overnight and secondary antibodies were incubated 1h at room temperature. The following antibodies were used: anti-HDAC6 Cell Signalling (7558), anti-Calnexin (E-10) (sc-46669), anti-Laminin (ab26300), anti-Histone H4 acetyl K12 (ab177793), anti-H4K16 ac Merck Millipore (07-329), anti-TH4 ac Merck Millipore (06-866), anti-H3k9 ac Merck Millipore (07-352) and anti-TH3 ac Merck Millipore (06-599), anti-Acetyl-α-tubulin (Cell Signaling), anti-α-tubulin (Calbiochem), anti-PARPc, anti-P53 and anti-Survivin Cell Signaling (2802), anti-β-actin Abcam (ab8227), secondary goat anti-mouse (Santa Cruz, St. Louis, MO, USA), anti-rabbit HRP Cell Signalling (7074) and anti-mouse HRP antibodies Cell Signalling (7076).

### MNase accessibility assay

Cells were lysed in NP-40 lysis buffer (ce-cold NP40 lysis buffer (10mM Tris [pH 7.4], 10mM NaCl, 3mM MgCl_2_, 0.5% NP-40, 0.15mM spermine, 0.5mM spermidine) and incubated on ice for 5minutes.). Nuclei were resuspended in Micrococal nuclease (MNase) digestion buffer (10mM Tris-HCl pH 8.0, 1mM CaCl_2_). A total of 0.06units of MNase Sigma-Aldrich (UK) was added to each sample and incubated at 15-20°C for 5minutes. The reaction was stopped by the addition of MNase digestion buffer, MNase stop buffer ((0.5 ml) - 5% SDS; 250mM EDTA), proteinase K and 20% SDS followed by overnight incubation. DNA was extracted by standard phenol/chloroform extraction and ethanol precipitation.

### Measurement of reactive oxygen species

HCT-116 cells were cultured for 24h and subsequently treated with different compounds over 2 hours: C1A (10μM), H_2_O_2_ (100μM) and/or N-acetylcystein (NAC from Sigma-Aldrich) (1mM). Cells were subsequently washed with PBS, replenished with phenol-red free medium with 10% serum supplemented with carboxy-H2DCFDA (C400-Life technologies) and incubated for 30min at 37°C. Optical densities were measured at 520nm with PHERAstar plate reader (BMG LABTECH Ltd, Aylesbury, UK).

### Comet assay

HCT-116 cells were treated for 3h with different concentration of 5μM of C1A or 3μM of Doxorubicin as positive controls. The assay was performed according to our previous study(65) from the original protocol of Singh et al.(66). Briefly, the standard slides were immersed vertically in 1% normal melting agarose (NMA) at 55°C and left vertically to allow the agarose to solidify. The slides were then kept at 4°C until use. Approximately 10,000 cells were mixed with 85μl of low-melting agarose (LMA; 0.7% in PBS) (FMC) at 37°C and, the cell suspension was rapidly pipetted onto the first agarose layer, spread using a coverslip and kept at 4°C for 8 min for the LMA to solidify. The coverslips were then removed, and a third layer of 100μl LMA (0.7%) at 37°C was added, covered with a coverslip, and again allowed to solidify at 4°C for 8 min. After the top layer of agarose was solidified, the slides were immersed in a chilled lysis solution made up of 2.5M NaCl, 0.1M Na2EDTA, 10-2M Tris–HCl, 1% sodium sarcosinate, pH 10, with 1% Triton X-100 and 10% DMSO added just before use. They were kept at 4°C in the dark for at least 1h to lyse the cells and to allow DNA unfolding. The slides were removed from the lysis solution, drained and placed on a horizontal gel electrophoresis unit, side by side. The tank was filled with chilled fresh alkaline solution (10-3M Na_2_ EDTA, 0.3M NaOH) at 4°C and pH 12.8, in order to detect double-and single-strand breaks as well as alkali-labile sites (67). Before electrophoresis, the slides were left in the solution for 20min to allow the unwinding of DNA. Electrophoresis was carried out at low temperature (4°C) for 20min at 1.6V cm-1 and 300mA. In order to prevent additional DNA damage, all the steps described above were conducted under yellow light or in the dark. After electrophoresis, slides were gently washed in a neutralization buffer (0.4M Tris–HCl, pH 7.5) to remove alkali and detergent, and stained with 50 l DAPI (5μg ml-1) in Vectashield (mounting medium for fluorescence H-1000, Vector Laboratories, USA). DNA of individual cells was viewed using an epifluorescence microscope OLYMPUS Vanox AHBT3, with an excitation filter of 550nm and barrier filter of 590nm, connected to a CCD camera and a pentium computer. Images of 50 randomly selected cells were captured by digitization from each sample. They were examined automatically using an image analysis CASys software (Synoptics Ltd., image processing systems, UK)(68). The measure of damage was tail moment, which is an integral of the distance and amount of DNA that has migrated out of the comet “head”. An increase of DNA tail moments over the control is a measure of DNA damage.

### Growth inhibitory assay

Drug concentrations that inhibited 50% of cell growth (GI50) were determined using a sulforhodamine B technique as described elsewhere (Vichai & Kirtikara, 2006).

### Cell Cycle

Cell flow cytometry analyses were conducted to evaluate cell cycle. HCT-116 cell line was exposed to 24h C1A drug treatment. Non-confluent cultures of exponentially growing cells were trypsinized and ethanol fixed. Cells lines were incubated in PBS containing propidium iodide and RNAse A from Invitrogen (12091021) for 2h after washes. Flow cytometry data was processed and analyzed with FlowJo software (Tree Star).

### Immunoprecipitation and Histone deacetylase activity assay

HCT-116 cells were washed 3 times with PBS and lysed with JLB containing a protease inhibitor cocktail (10μg/mL aprotinin, 10μg/mL leupeptin, 1mM PMSF) at 4°C. The HDAC6 antibody was incubated with cell extract for 2h at 4°C on a rotating platform. Immunocomplexes were collected using protein A/G agarose beads (Santa Cruz Biotechnology). The beads were collected by centrifugation, and washed first with JLB, and then with histone deacetylase buffer (Upstate Cell Signaling Solutions). The histone deacetylase assay was performed using the fluorometric HDAC assay kit (Upstate Cell Signaling Solutions) as per the manufacturer’s recommendations. The immunoprecipitates from each reaction were incubated with 100μM substrate and test compounds for 1hour at 37°C. Fluorescence was read using a Millipore Cytofluor 2300 Fluorescence Plate Reader (Millipore, Billerica, MA).

### Acetylome

HCT-116 cells were cultured for 24h and subsequently treated for 3h with C1A at 10μM. Protein samples were subsequently prepared in Pierce IP lysis buffer Thermo Fisher Scientific (Waltham, MA, USA) supplemented with protease and phosphatase inhibitor cocktails (Sigma) immunoprecipitated using Dynabeads® M-280 according to manufacturer’s instructions (Life technologies). Briefly, protein samples were incubated with pre-cleared beads coupled with acetyl-lysine antibody (Santa Cruz) for 1h on rotor at 4°C, washed and the supernatant removed using a magnetic stand (Millipore). NanoLC-MS/MS was performed by Applied Biomics, Inc (Hayward, CA, USA). Proteins were eluted from beads and supernatant was concentrated using 5K MW cut off spin columns. Proteins were then exchanged into 50mM ammonium bicarbonate buffer. DTT was added to a final concentration of 10mM and incubated at 60°C for 30min, followed by cooling down to RT. Iodoacetamide was then added to a final concentration of 10mM and incubated in the dark for 30min at RT. The proteins were then digested by trypsin Promega (Madison, WI, USA) overnight at 37°C. NanoLC was carried out using a Dionex Ultimate 3000 Milford (MA, USA). Fractions were collected at 20second intervals followed by Mass Spectrometry analysis on AB SCIEX TOF/TOFTM 5800 System (AB SCIEX). Both of the resulting peptide mass and the associated fragmentation spectra were submitted to GPS Explorer workstation equipped with MASCOT search engine MatrixScience (London, UK) to search the Swiss Prot database. Searches were performed without constraining protein MW or isoelectric point, with variable carbamidomethylation of cysteine and oxidation of methionine residues, and with one missed cleavage.

### RNA Extraction and Real-Time RTqPCR

RNA was isolated from cell lines using miRVana miRNA Isolation Kit (Ambion; Life Technologies, USA). The quantity and quality of the total RNA was determined with Nanodrop ND-2000 Spectrophotometer (Thermo Scientific). Prior reverse transcription was performed using TaqMan Reverse Transcription Kit (Applied Biosystems; Life Technologies) in GeneAmp PCR 9700 thermocycler and qRT-PCR amplification with TaqMan Universal PCR Master Mix (Applied Biosystems). All qRT-PCR measurements were obtained in a 7900HT Fast Real Time PCR System with ExpressionSuite Software v1.0 (Applied Biosystems). Taqman probes utilized in this study are listed in S-Table 9.

### Patients and clinical samples

This study includes two independent series of colorectal samples obtained between 1993 and 2016. Series 1 comprised 7 paired samples (normal and tumor tissues) frozen samples. Series 2 comprised a tissue microarray (TMA) with 113 paraffin-embedded samples. This series consisted of 51 paired tumor and normal tissues, and 11 tumor samples with no matched normal tissues. Rhabdomyosarcoma samples patient comprised a TMA with 13 paired tumor and adjacent normal tissues. Samples were obtained between 2012 and 2016. Glioblastoma samples patient comprised a TMA with 13 paired tumor and adjacent normal tissues. Samples were obtained between 2009 and 2016. The patient characteristics of colorectal series 2, rhabdomyosarcoma and gliobastoma samples are summarized in S-Table 4.

Colorectal series 2, rhabdomyosarcoma and gliobastoma samples was obtained from the Department of Pathology at the Hospital Universitario Virgen del Rocío (Seville, Spain) and HUVR-IBiS Biobank. Approval of the Ethics Committee of our institution was obtained as well as the written informed consent before including samples and data in the HUVR-IBiS Biobank as provided in law 14/2007, 3 July, on Biomedical Research.

### Immunohistochemistry HDAC6

Five-micrometer-thick tissue sections from tissue microarray blocks were dewaxed in xylene and rehydrated in a series of graded alcohols. Sections were immersed in 3% H_2_O_2_ aqueous solution for 30min to exhaust endogenous peroxidase activity and then covered with 1% blocking reagent Roche (Mannheim, Germany) in 0.05% Tween 20-PBS, to block nonspecific binding sites. Antigen retrieval was performed with a pressure cooker, using EDTA buffer, pH 9.0 Dako (Glostrup, Denmark). Sections were incubated with primary antibodies (HDAC 1/600, Cell Signaling (7558)) overnight at 4°C. Peroxidase-labeled secondary antibodies and 3,3-diaminobenzidine were applied to develop immunoreactivity, according to manufacturer’s protocol EnVision (Dako). Slides were then counterstained with hematoxylin and mounted in DPX BDH Laboratories (Poole, UK). Sections in which primary antibody was omitted were used as negative controls. Immunostaining was evaluated independently by two observers.

### Tissue microarray (TMA)

Tissue sections (5μm) from formalin-fixed paraffin-embedded (FFPE) of colorectal cancer, rhabdomyosarcoma and gliobastoma were stained with haematoxylin and eosin. Representative malignant areas from samples were carefully selected from the stained sections of each tumor, and two 1mm diameter tissue cores were obtained from each sample to build up the TMA in duplicate.

### Tumor xenografts in mice

The in vivo study was approved by The University of Seville Ethical Committee for Experimental Research and fulfilled the requirements for experimental animal research in accordance the Spanish regulations (BOE 34/11370-421, 2013) for the use of laboratory animals. After the approval of the Institutional Animal Research Ethics Committee, 10 Balb/c nude mice (Harlan) were injected subcutaneously with 4.5×10^6^ HTC-116 cells in a 1:1 proportion (DMEM medium and Matrigel Matrix (Becton Dickinson)). Balb/c nude mice bearing 100-400mm^3^ tumors in one flank were randomized in 2 groups. One group received an intraperitoneal injection (IP) of 40mg/kg C1A once daily for two weeks and a second group was treated with the vehicle (saline) under the same regimen (control group). Tumor growth was measured with a digital caliper and mice were sacrificed when tumor volumes reached tolerable size limits and animals were euthanized by anesthetic overdose. All animals were housed and handled in accordance with the Guide for the Care and Use of Laboratory Animals. Tumor response was evaluated and compared between both groups. At the end of the experiment tumors were excised and frozen in liquid nitrogen.

### Gene array

Total RNA was extracted from mice tumors using RNeasy mini kit (Qiagen (Crawley, UK) and hybridised to Affymetrix human genome U219 microarray Affymetrix (Santa Clara, CA, USA). Studies were performed under contract by AlphaMetrix biotech (Rödermark, Germany). A differential expression of 1.5-fold was selected as a cut-off.

### Mitochondrial membrane depolarisation and cell death assessment

Cells were trypsinised, washed, and re-suspended in PBS 1X and distributed in flow cytometry tubes. Each tube contains 150μl of PBS and 40nM 3,3-dihexaoxacarbocyanine iodide (DiOC6) for measuring mitochondrial membrane depolarisation loss. Cells viability was assessed by staining with Propidium iodide (PI) (50μM) for 40minutes at 37°C. Cells were then analyzed using FACSCalibur TM (BD Biosciences) machine with the cell-Quest program.

### Statistical analysis

Differential expression between two groups was evaluated with the T-student test, and with the 1 way Anova test for more than two groups, followed by Bonferroni multiple comparison post-test. Paired samples t-test was applied to compare the mean level of expression within the same specimens. Fisher’s exact test was used to evaluate differences between HDAC6 expression and the present or absence of metastasis in patients. Overall survival and disease-free survival were analyzed using the Kaplan-Meier estimator, and the differences were evaluated using the log-rank test. Cox regression proportional hazards models were used for estimation of hazard ratios (HRs) for disease free survival from colorectal tumor patients in both uni- and multivariable analysis, adjusted for HDAC6 expression and clinical stage.

For all analyses, data represent mean±standard deviation (s.d).; p-values of ≤0.05 were considered statistically significant. Univariate analyses were performed using the Prism 4.0 software (GraphPad) and Cox regression analyses were performed using SPSS software version 22 (IBM, Armonk, NY, USA).

## References

1. Aldana-Masangkay GI, and Sakamoto KM. The role of HDAC6 in cancer. Journal of biomedicine & biotechnology. 2011;2011:875824.

2. Bertos NR, Gilquin B, Chan GK, Yen TJ, Khochbin S, and Yang XJ. Role of the tetradecapeptide repeat domain of human histone deacetylase 6 in cytoplasmic retention. J Biol Chem. 2004;279(46):48246–54.

3. Boyault C, Zhang Y, Fritah S, Caron C, Gilquin B, Kwon SH, et al. HDAC6 controls major cell response pathways to cytotoxic accumulation of protein aggregates. Genes Dev. 2007;21(17):2172–81.

4. Fernandes I, Bastien Y, Wai T, Nygard K, Lin R, Cormier O, et al. Ligand-dependent nuclear receptor corepressor LCoR functions by histone deacetylase-dependent and - independent mechanisms. Mol Cell. 2003;11(1):139–50.

5. Glasgow JE, and Daniele RP. Role of microtubules in random cell migration: stabilization of cell polarity. Cell motility and the cytoskeleton. 1994;27(1):88–96.

6. Ryu HW, Lee DH, Shin DH, Kim SH, and Kwon SH. Aceroside VIII is a new natural selective HDAC6 inhibitor that synergistically enhances the anticancer activity of HDAC inhibitor in HT29 cells. Planta medica. 2015;81(3):222–7.

7. Lee HY, Tsai AC, Chen MC, Shen PJ, Cheng YC, Kuo CC, et al. Azaindolylsulfonamides, with a more selective inhibitory effect on histone deacetylase 6 activity, exhibit antitumor activity in colorectal cancer HCT116 cells. J Med Chem. 2014;57(10):4009–22.

8. Yang Z, Wang T, Wang F, Niu T, Liu Z, Chen X, et al. Discovery of Selective Histone Deacetylase 6 Inhibitors Using the Quinazoline as the Cap for the Treatment of Cancer. J Med Chem. 2016;59(4):1455–70.

9. Chao OS, Chang TC, Di Bella MA, Alessandro R, Anzanello F, Rappa G, et al. The HDAC6 Inhibitor Tubacin Induces Release of CD133(+) Extracellular Vesicles From Cancer Cells. J Cell Biochem. 2017;118(12):4414–24.

10. Gotze S, Coersmeyer M, Muller O, and Sievers S. Histone deacetylase inhibitors induce attenuation of Wnt signaling and TCF7L2 depletion in colorectal carcinoma cells. Int J Oncol. 2014;45(4):1715–23.

11. Lee SJ, Li Z, Litan A, Yoo S, and Langhans SA. EGF-induced sodium influx regulates EGFR trafficking through HDAC6 and tubulin acetylation. BMC cell biology. 2015;16:24.

12. Kaliszczak M, van Hechanova E, Li Y, Alsadah H, Parzych K, Auner HW, et al. The HDAC6 inhibitor C1A modulates autophagy substrates in diverse cancer cells and induces cell death. Br J Cancer. 2018;119(10):1278–87.

13. Zhang Y, Kwon S, Yamaguchi T, Cubizolles F, Rousseaux S, Kneissel M, et al. Mice lacking histone deacetylase 6 have hyperacetylated tubulin but are viable and develop normally. Mol Cell Biol. 2008;28(5):1688–701.

14. Lee YS, Lim KH, Guo X, Kawaguchi Y, Gao Y, Barrientos T, et al. The cytoplasmic deacetylase HDAC6 is required for efficient oncogenic tumorigenesis. Cancer Res. 2008;68(18):7561–9.

15. Kaluza D, Kroll J, Gesierich S, Yao TP, Boon RA, Hergenreider E, et al. Class IIb HDAC6 regulates endothelial cell migration and angiogenesis by deacetylation of cortactin. EMBO J. 2011;30(20):4142–56.

16. Haggarty SJ, Koeller KM, Wong JC, Grozinger CM, and Schreiber SL. Domain-selective small-molecule inhibitor of histone deacetylase 6 (HDAC6)-mediated tubulin deacetylation. Proc Natl Acad Sci U S A. 2003;100(8):4389–94.

17. Parmigiani RB, Xu WS, Venta-Perez G, Erdjument-Bromage H, Yaneva M, Tempst P, et al. HDAC6 is a specific deacetylase of peroxiredoxins and is involved in redox regulation. Proc Natl Acad Sci U S A. 2008;105(28):9633–8.

18. Zhang X, Yuan Z, Zhang Y, Yong S, Salas-Burgos A, Koomen J, et al. HDAC6 modulates cell motility by altering the acetylation level of cortactin. Mol Cell. 2007;27(2):197–213.

19. Witt O, Deubzer HE, Milde T, and Oehme I. HDAC family: What are the cancer relevant targets? Cancer Lett. 2009;277(1):8–21.

20. Bali P, Pranpat M, Bradner J, Balasis M, Fiskus W, Guo F, et al. Inhibition of histone deacetylase 6 acetylates and disrupts the chaperone function of heat shock protein 90: a novel basis for antileukemia activity of histone deacetylase inhibitors. J Biol Chem. 2005;280(29):26729–34.

21. Butler KV, Kalin J, Brochier C, Vistoli G, Langley B, and Kozikowski AP. Rational design and simple chemistry yield a superior, neuroprotective HDAC6 inhibitor, tubastatin A. Journal of the American Chemical Society. 2010;132(31):10842–6.

22. Kaliszczak M, Trousil S, Aberg O, Perumal M, Nguyen QD, and Aboagye EO. A novel small molecule hydroxamate preferentially inhibits HDAC6 activity and tumour growth. Br J Cancer. 2013;108(2):342–50.

23. Suraweera A, O’Byrne KJ, and Richard DJ. Combination Therapy With Histone Deacetylase Inhibitors (HDACi) for the Treatment of Cancer: Achieving the Full Therapeutic Potential of HDACi. Front Oncol. 2018;8:92.

24. Lee DH, Won HR, Ryu HW, Han JM, and Kwon SH. The HDAC6 inhibitor ACY1215 enhances the anticancer activity of oxaliplatin in colorectal cancer cells. Int J Oncol. 2018;53(2):844–54.

25. Lachmann A, Torre D, Keenan AB, Jagodnik KM, Lee HJ, Wang L, et al. Massive mining of publicly available RNA-seq data from human and mouse. Nat Commun. 2018;9(1):1366.

26. Ashktorab H, Belgrave K, Hosseinkhah F, Brim H, Nouraie M, Takkikto M, et al. Global histone H4 acetylation and HDAC2 expression in colon adenoma and carcinoma. Dig Dis Sci. 2009;54(10):2109–17.

27. Alzoubi S, Brody L, Rahman S, Mahul-Mellier AL, Mercado N, Ito K, et al. Synergy between histone deacetylase inhibitors and DNA-damaging agents is mediated by histone deacetylase 2 in colorectal cancer. Oncotarget. 2016;7(28):44505–21.

28. Katan-Khaykovich Y, and Struhl K. Dynamics of global histone acetylation and deacetylation in vivo: rapid restoration of normal histone acetylation status upon removal of activators and repressors. Genes Dev. 2002;16(6):743–52.

29. Boffa LC, Walker J, Chen TA, Sterner R, Mariani MR, and Allfrey VG. Factors affecting nucleosome structure in transcriptionally active chromatin. Histone acetylation, nascent RNA and inhibitors of RNA synthesis. Eur J Biochem. 1990;194(3):811–23.

30. McCool KW, Xu X, Singer DB, Murdoch FE, and Fritsch MK. The role of histone acetylation in regulating early gene expression patterns during early embryonic stem cell differentiation. J Biol Chem. 2007;282(9):6696–706.

31. Kao J, Salari K, Bocanegra M, Choi YL, Girard L, Gandhi J, et al. Molecular profiling of breast cancer cell lines defines relevant tumor models and provides a resource for cancer gene discovery. PLoS One. 2009;4(7):e6146.

32. Borkar SA, Lakshmiprasad G, Subbarao KC, Sharma MC, and Mahapatra AK. Giant cell glioblastoma in the pediatric age group: Report of two cases. Journal of pediatric neurosciences. 2013;8(1):38–40.

33. Fu W, Asp P, Canter B, and Dynlacht BD. Primary cilia control hedgehog signaling during muscle differentiation and are deregulated in rhabdomyosarcoma. Proc Natl Acad Sci U S A. 2014;111(25):9151–6.

34. Liu H, Zang C, Fenner MH, Possinger K, and Elstner E. PPARgamma ligands and ATRA inhibit the invasion of human breast cancer cells in vitro. Breast cancer research and treatment. 2003;79(1):63–74.

35. Chavez KJ, Garimella SV, and Lipkowitz S. Triple negative breast cancer cell lines: one tool in the search for better treatment of triple negative breast cancer. Breast disease. 2010;32(1–2):35–48.

36. Lee EJ, Lee HG, Park SH, Choi EY, and Park SH. CD99 type II is a determining factor for the differentiation of primitive neuroectodermal cells. Experimental & molecular medicine. 2003;35(5):438–47.

37. Snoek R, Bruchovsky N, Kasper S, Matusik RJ, Gleave M, Sato N, et al. Differential transactivation by the androgen receptor in prostate cancer cells. Prostate. 1998;36(4):256–63.

38. Hai Y, and Christianson DW. Histone deacetylase 6 structure and molecular basis of catalysis and inhibition. Nature chemical biology. 2016;12(9):741–7.

39. Zhang SL, Zhu HY, Zhou BY, Chu Y, Huo JR, Tan YY, et al. Histone deacetylase 6 is overexpressed and promotes tumor growth of colon cancer through regulation of the MAPK/ERK signal pathway. Onco Targets Ther. 2019;12:2409–19.

40. Sixto-Lopez Y, Gomez-Vidal JA, de Pedro N, Bello M, Rosales-Hernandez MC, and Correa-Basurto J. Hydroxamic acid derivatives as HDAC1, HDAC6 and HDAC8 inhibitors with antiproliferative activity in cancer cell lines. Sci Rep. 2020;10(1):10462.

41. Scholz C, Weinert BT, Wagner SA, Beli P, Miyake Y, Qi J, et al. Acetylation site specificities of lysine deacetylase inhibitors in human cells. Nat Biotechnol. 2015;33(4):415–23.

42. Song H, Li CW, Labaff AM, Lim SO, Li LY, Kan SF, et al. Acetylation of EGF receptor contributes to tumor cell resistance to histone deacetylase inhibitors. Biochem Biophys Res Commun. 2011;404(1):68–73.

43. Lai CJ, Bao R, Tao X, Wang J, Atoyan R, Qu H, et al. CUDC-101, a multitargeted inhibitor of histone deacetylase, epidermal growth factor receptor, and human epidermal growth factor receptor 2, exerts potent anticancer activity. Cancer Res. 2010;70(9):3647–56.

44. He L, Gao L, Shay C, Lang L, Lv F, and Teng Y. Histone deacetylase inhibitors suppress aggressiveness of head and neck squamous cell carcinoma via histone acetylation-independent blockade of the EGFR-Arf1 axis. Journal of experimental & clinical cancer research: CR. 2019;38(1):84.

45. Tu CY, Chen CH, Hsia TC, Hsu MH, Wei YL, Yu MC, et al. Trichostatin A suppresses EGFR expression through induction of microRNA-7 in an HDAC-independent manner in lapatinib-treated cells. Biomed Res Int. 2014;2014:168949.

46. Yokota T. Are KRAS/BRAF mutations potent prognostic and/or predictive biomarkers in colorectal cancers? Anti-cancer agents in medicinal chemistry. 2012;12(2):163–71.

47. Yokota T, Ura T, Shibata N, Takahari D, Shitara K, Nomura M, et al. BRAF mutation is a powerful prognostic factor in advanced and recurrent colorectal cancer. Br J Cancer. 2011;104(5):856–62.

48. Alberts SR, Sargent DJ, Nair S, Mahoney MR, Mooney M, Thibodeau SN, et al. Effect of oxaliplatin, fluorouracil, and leucovorin with or without cetuximab on survival among patients with resected stage III colon cancer: a randomized trial. Jama. 2012;307(13):1383–93.

49. Namdar M, Perez G, Ngo L, and Marks PA. Selective inhibition of histone deacetylase 6 (HDAC6) induces DNA damage and sensitizes transformed cells to anticancer agents. Proc Natl Acad Sci U S A. 2010;107(46):20003–8.

50. Haberland M, Montgomery RL, and Olson EN. The many roles of histone deacetylases in development and physiology: implications for disease and therapy. Nat Rev Genet. 2009;10(1):32–42.

51. Ran J, Yang Y, Li D, Liu M, and Zhou J. Deacetylation of alpha-tubulin and cortactin is required for HDAC6 to trigger ciliary disassembly. Sci Rep. 2015;5:12917.

52. Li Y, Shin D, and Kwon SH. Histone deacetylase 6 plays a role as a distinct regulator of diverse cellular processes. The FEBS journal. 2013;280(3):775–93.

53. Liu Y, Peng L, Seto E, Huang S, and Qiu Y. Modulation of histone deacetylase 6 (HDAC6) nuclear import and tubulin deacetylase activity through acetylation. J Biol Chem. 2012;287(34):29168–74.

54. Asthana J, Kapoor S, Mohan R, and Panda D. Inhibition of HDAC6 deacetylase activity increases its binding with microtubules and suppresses microtubule dynamic instability in MCF-7 cells. J Biol Chem. 2013;288(31):22516–26.

55. Sigismund S, Avanzato D, and Lanzetti L. Emerging functions of the EGFR in cancer. Molecular oncology. 2018;12(1):3–20.

56. Lu Z, Ghosh S, Wang Z, and Hunter T. Downregulation of caveolin-1 function by EGF leads to the loss of E-cadherin, increased transcriptional activity of beta-catenin, and enhanced tumor cell invasion. Cancer Cell. 2003;4(6):499–515.

57. Li Y, and Seto E. HDACs and HDAC Inhibitors in Cancer Development and Therapy. Cold Spring Harbor perspectives in medicine. 2016;6(10).

58. Muhammad S, Jiang Z, Liu Z, Kaur K, and Wang X. The role of EGFR monoclonal antibodies (MoABs) cetuximab/panitumab, and BRAF inhibitors in BRAF mutated colorectal cancer. Journal of gastrointestinal oncology. 2013;4(1):72–81.

59. Li T, Zhang C, Hassan S, Liu X, Song F, Chen K, et al. Histone deacetylase 6 in cancer. Journal of hematology & oncology. 2018;11(1):111.

60. Tambunan US, Bramantya N, and Parikesit AA. In silico modification of suberoylanilide hydroxamic acid (SAHA) as potential inhibitor for class II histone deacetylase (HDAC). BMC Bioinformatics. 2011;12 Suppl 13:S23.

61. Qiu L, Burgess A, Fairlie DP, Leonard H, Parsons PG, and Gabrielli BG. Histone deacetylase inhibitors trigger a G2 checkpoint in normal cells that is defective in tumor cells. Mol Biol Cell. 2000;11(6):2069–83.

62. Wu YW, Hsu KC, Lee HY, Huang TC, Lin TE, Chen YL, et al. A Novel Dual HDAC6 and Tubulin Inhibitor, MPT0B451, Displays Anti-tumor Ability in Human Cancer Cells in Vitro and in Vivo. Front Pharmacol. 2018;9:205.

63. Hajji N, Wallenborg K, Vlachos P, Fullgrabe J, Hermanson O, and Joseph B. Opposing effects of hMOF and SIRT1 on H4K16 acetylation and the sensitivity to the topoisomerase II inhibitor etoposide. Oncogene. 2010;29(15):2192–204.

64. Shechter D, Dormann HL, Allis CD, and Hake SB. Extraction, purification and analysis of histones. Nat Protoc. 2007;2(6):1445–57.

65. Hajji N, Pastor N, Mateos S, Dominguez I, and Cortes F. DNA strand breaks induced by the anti-topoisomerase II bis-dioxopiperazine ICRF-193. Mutat Res. 2003;530(1–2):35–46.

66. Singh NP, McCoy MT, Tice RR, and Schneider EL. A simple technique for quantitation of low levels of DNA damage in individual cells. Exp Cell Res. 1988;175(1):184–91.

67. Fairbairn DW, Olive PL, and O’Neill KL. The comet assay: a comprehensive review. Mutat Res. 1995;339(1):37–59.

68. Olive PL, Banath JP, and Durand RE. Detection of etoposide resistance by measuring DNA damage in individual Chinese hamster cells. J Natl Cancer Inst. 1990;82(9):779–83.

