## supplementary data. Dani et al HDAC6 for "Novel nuclear role of HDAC6 in prognosis and therapeutic target for colorectal cancer"

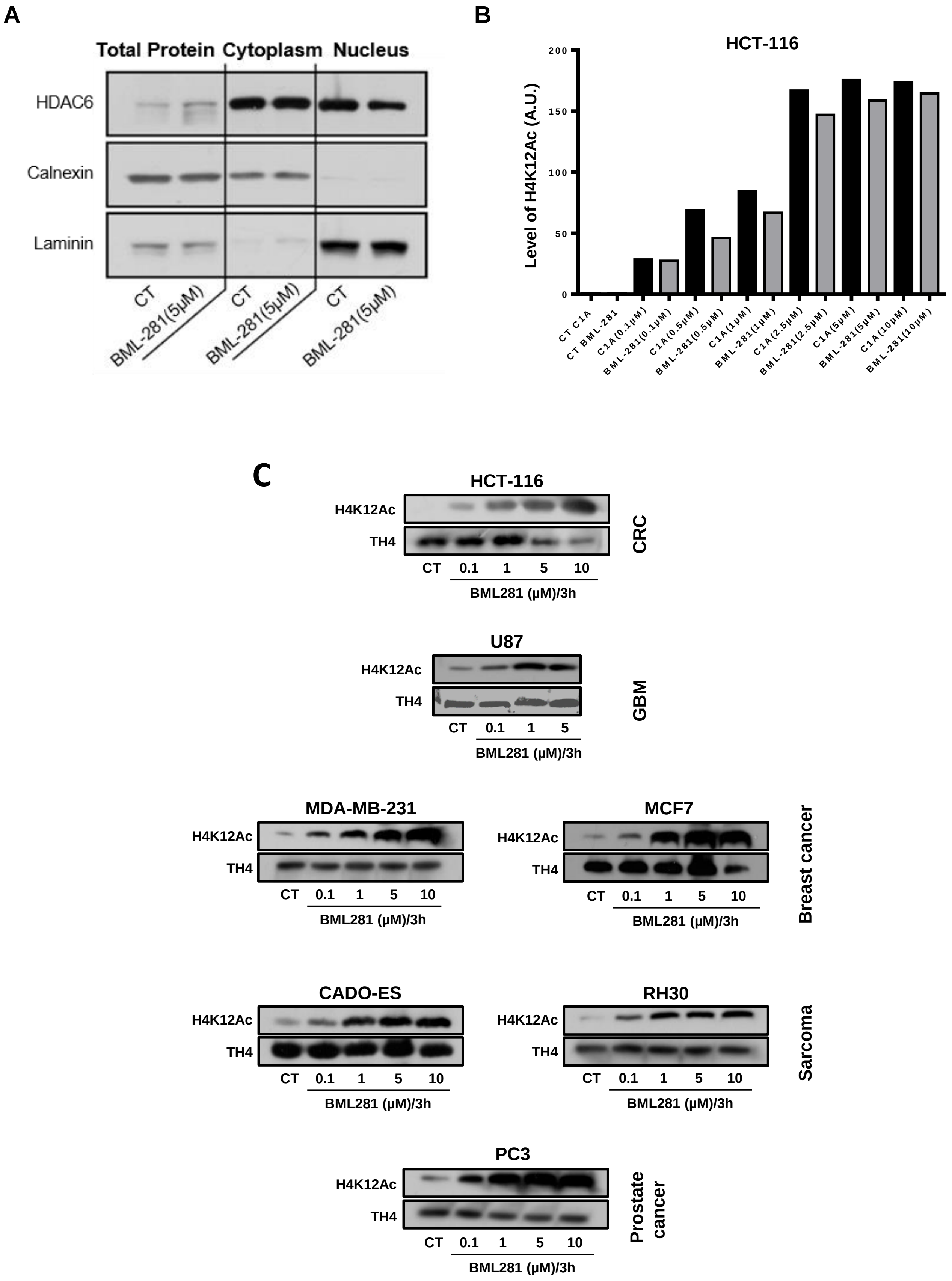


### Supplementary Figure1: S1

**Supplementary Figure Legends**

**Supplementary Figure 1: S1. HDAC6 regulates the level of H4K12ac in distinct cancer cell types.**

Assessment of HDAC6 localization in cytoplasmic and nuclear fractions. HCT-116 cells were treated with BML-281, 5µM for 3h and both fractions cytoplasmic and nuclear were subjected to immunoblotting (A). Measurement of H4K12 acetylation level after selective inhibition of HDAC6 by C1A or BML-281 in HCT-116 cells (B). Assessment of H4K12ac level in different cancer types after HDAC6 inhibition by BML-281 (C).

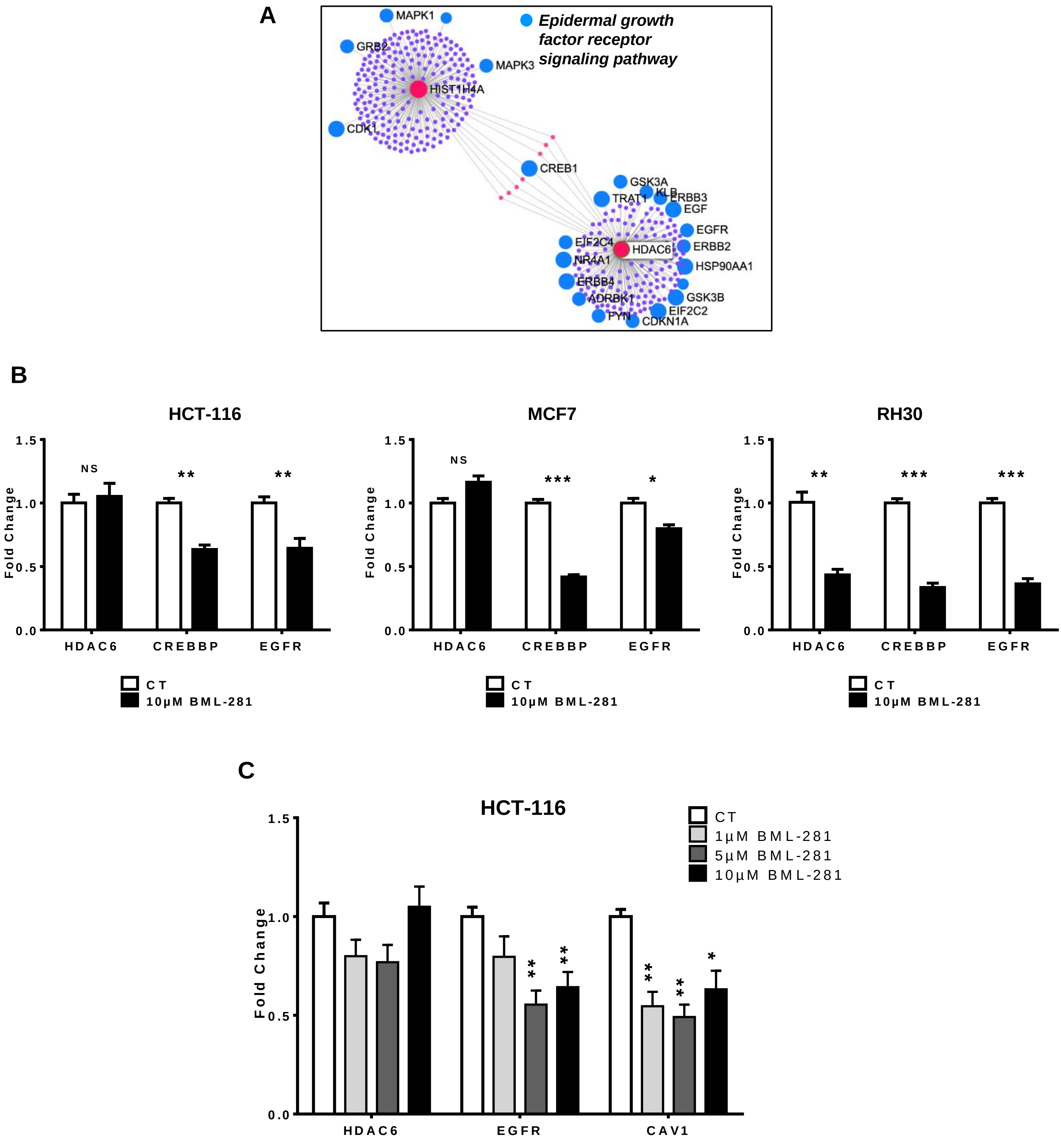


### Supplementary Figure 2: S2

**Supplementary Figure 2: S2.**

Identification of EGFR signaling pathway on predicted protein-protein interaction between HDAC6–HIST1H4E analyzed by networkanalyst using reactome data base (A). QPCR measurement of *EGFR*, *CREBBP* and *HDAC6* expression level *in* HCT-116, MCF7 and RH30 cell lines treated with 10µM of BML-281 for 3h (B). Assessment of *HDAC6*, *EGFR* and *CAV1* expression in HCT-116 treated with varying concentrations of BML-281. 2-way ANOVA: p-value<0.05(*), p-value<0.01(**), p-value<0.001(***).

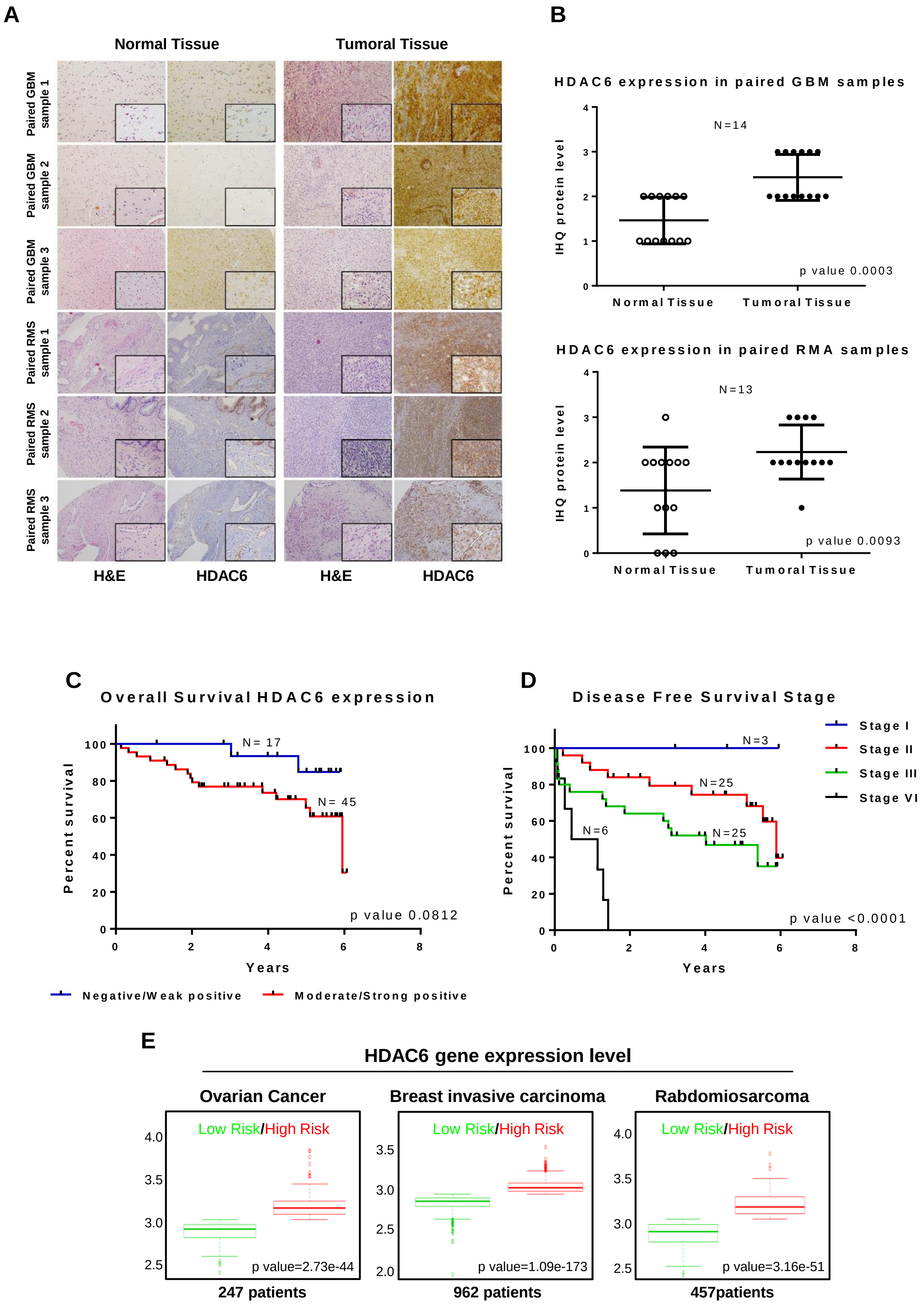


**Supplementary Figure 3: S3**

**Supplementary Figure 3: S3.**

IHQ representative images and quantification of the expression of HDAC6 in Glioblastoma (GBM) and Rhabdomyosarcoma (RMS) patients (paired tumor and adjacent normal tissues) (A and B). Kaplan–Meier of overall survival curves based on the HDAC6 expression by IHC in colorectal cancer (CRC) patients (C). Kaplan–Meier representation of the association of CRC clinical stages with DFS (D). TCGA data analysis of risk factor correlation with HDAC6 expression in ovarian, breast and sarcoma cancer patients using survExpress website "http://bioinformatica.mty.itesm.mx/SurvExpress" (E).

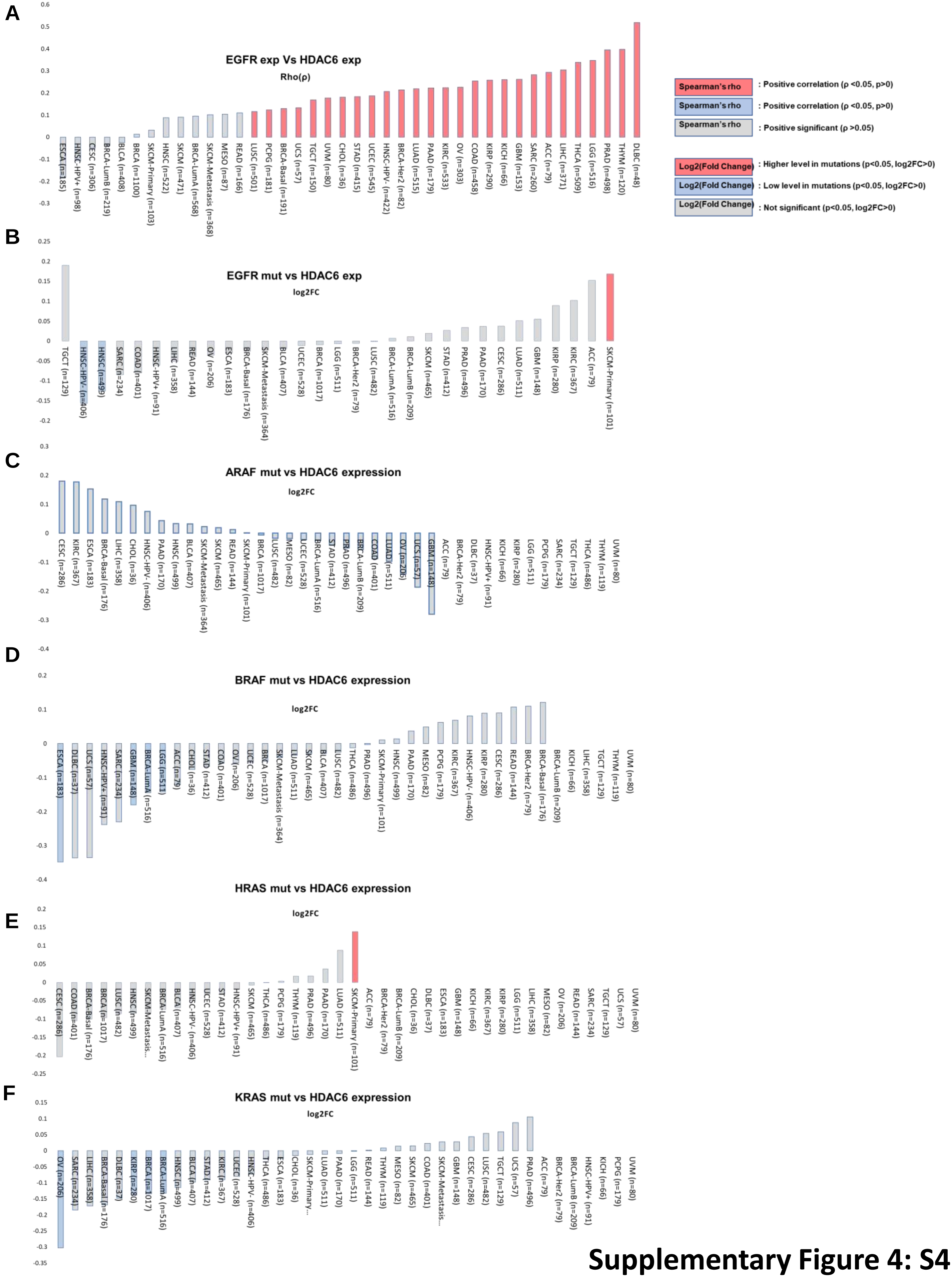


**Supplementary Figure 4: S4**

**Supplementary Figure 4: S4**

TCGA data analysis of 40 cancers types and representation of the correlation between key colorectal cancer driver mutations and HDAC6 expression. Correlation analysis between *EGFR expression* and *HDAC6* expression (A). Correlation analysis between mutated *EGFR* and *HDAC6* expression (B).

Correlation analysis between HDAC6 expression and KRAS*, HRAS, ARAF* and *BRAF* mutations (C-F). TCGA data analysis was based on the Gene_DE module to study the differential expression between tumor and adjacent normal tissues for gene of interest across all 40 types of cancer (http://timer.cistrome.org/).
